# A neurovascular progenitor sits at the nexus of glioblastoma lineage trajectories

**DOI:** 10.1101/2024.07.24.604840

**Authors:** Elisa Fazzari, Daria J. Azizad, Kwanha Yu, Weihong Ge, Matthew X. Li, Patricia R. Nano, Shivani Baisiwala, Antoni Martija, Ryan L. Kan, Jaela Caston, Loukas N. Diafos, Hong A. Tum, Christopher Tse, Nicholas A. Bayley, Vjola Haka, Dimitri Cadet, Travis Perryman, Jose A. Soto, Brittney Wick, Avinash Veerappa, Chittibabu Guda, Lauren W. Powers, Vaibhav Jain, Michael Aksu, Richard G. Everson, Marvin Bergsneider, David R. Raleigh, Simon G. Gregory, Elizabeth E. Crouch, Kunal S. Patel, Linda M. Liau, David A. Nathanson, Benjamin Deneen, Aparna Bhaduri

## Abstract

Glioblastoma (GBM) exhibits developmental programs and marked cellular heterogeneity, yet how these features are organized into connected lineage hierarchies remains unclear. Here we identify a rare tumor-intrinsic population, termed the neurovascular progenitor (NVP), that occupies an intermediate position between the major GBM organizational axes. NVP cells co-express neural progenitor and perivascular transcriptional features, are consistently detected across independent patient cohorts, retain canonical GBM copy-number alterations, and localize *in situ* in both vessel-associated and parenchymal niches. Using direct-from-patient lineage tracing in a human organoid tumor transplantation system, we show that individual NVP cells clonally generate both neural-like and mesenchymal/vascular-like malignant progeny, providing a concrete lineage link between states that are commonly considered mutually exclusive. Despite comprising ∼1% of tumor cells, NVP-derived lineages account for a majority of observed tumor cell types and disproportionately contribute to cycling compartments. Orthogonally, ablation of NVP-associated programs in an *in vivo* GBM model remodels tumor composition, elicits compensatory progenitor states, and significantly prolongs survival. Together, these findings position NVP as a fate-restricted yet highly influential lineage intermediate that serves as a functional bridge and organizational nexus within GBM hierarchies, linking population-level lineage architecture to the behavior of a specific progenitor cell type.

## Introduction

Glioblastoma (GBM) is the most common and aggressive form of primary adult brain cancer, carrying a dismal prognosis with a median survival of about 21 months following diagnosis^1^. Standard of care therapy largely employs nonspecific treatments including radiation and temozolomide chemotherapy^2,3^ that inevitably result in tumor recurrence. Extensive molecular and spatial characterization has shown that GBM displays immense inter- and intra-tumoral heterogeneity^4–18^. Previous studies indicate that there exist transcriptionally distinct cancer stem cell-like populations within GBM that may map to functionally distinct progenitor populations, similar to the diversity observed in human brain development^4,14^. For example, we have previously shown outer radial glia (oRG) to be a discrete cell type with both transcriptional and cell behavioral features that drives tumor migration^4^. This idea of what we refer to here as “progenitor cell types” is not at odds with previous models showing a shift between cell states within the tumor^10^ and instead establishes a framework for which additional experimentation is necessary to characterize the diversity of progenitors within the tumor and each of their functional roles. While much work has been done to postulate on the tumor-initiating cells of GBM, it is also important to understand the convergence of these lineages at the time of disease in order to predict how the mature tumor will evolve based on its fate-restricted cell types. Thus, beyond defining tumor-initiating cells, it is essential to understand how intermediate progenitor compartments sustain lineage output within established tumors, as these regenerative intermediates may preserve heterogeneity even if upstream populations are eliminated.

Understanding the landscape of progenitor subtypes within GBM patient tumors offers a window to understand the ability for GBM to evolve and adapt, as we interrogate in this study. Open questions remain regarding the functional role of GBM tumor heterogeneity, including granular descriptions of what progenitor populations exist and their unique roles within the tumor.

To characterize the landscape of progenitor cell types in human primary GBM, we compiled a meta-atlas of single-cell transcriptomes from seven published studies^4,6–8,10,16,17^. In addition to the transcriptomic landscape of GBM, we aimed to map the function of these cell types. In a companion study^19^, referred to here as the “CellTagging” dataset, we sought to functionally interrogate the diverse progenitors in GBM using a DNA barcoding approach in human GBM samples transplanted onto human cortical organoids. This study described the wide landscape of progenitor populations and described their varying likelihoods to contribute to tumor propagation.

One previously undescribed progenitor stood out from both the meta-atlas and CellTagging datasets. This progenitor cell co-expressed transcriptional programs previously thought to be associated with vascular or perivascular populations, so we named this population NVP, or neurovascular progenitor. WIth a transcriptional profile similar but distinct in key ways from radial glia, we classified this progenitor as its own cell type based on its distinct clone profile to other progenitors in the dataset. Not only was its transcriptional profile distinct, but of all progenitors, NVP had the highest functional entropy in the tumor, giving rise to the highest diversity of cell types. Here, we validate the existence of NVP in seven combined cross-institutional datasets, one recently published large scale GBM dataset which is stringently filtered for tumor cells and characterized GBM expression programs^20^, and ultimately *in situ*. In each, we observed the NVP cell population and sought to characterize it functionally as a paradigm of progenitor-specific behavior in GBM. We describe their morphological and molecular features, and use both *in vivo* and direct-from-patient systems to characterize their function. We demonstrate that NVP cells have dual-fate capabilities in clonally generating neuron-like and mesenchymal tumor populations, and that they generate a defined fraction of the overall tumor. Despite their low abundance in patient tumors, they disproportionately generate cycling cells and are competent to give rise to the majority, but not all, of tumor cell subtypes. Furthermore, elimination of NVP from an *in vivo* tumor model results in extended survival, highlighting the functional relevance of this clonal role within the tumor.

## Results

### Generation of a GBM-specific meta-atlas discovery of the NVP cell population

Previous efforts have transcriptionally profiled GBM at a single-cell level, and we reasoned that compiling these datasets could allow us to uncover novel progenitor populations essential to driving the tumor. Thus, we generated a meta-atlas of existing single-cell RNA sequencing (scRNA-seq) datasets^4,6,8,10,16,17,20^. While many of these datasets have been previously analyzed and aggregated, our approach of including only the IDH1 wild-type (IDH1^WT^), adult, direct-from-patient samples accounts for the unique nature of GBM compared to other gliomas and removes any contribution of *in vitro* culture methods, cell type selection, or distinct pathologies that may interfere with cell type identification. We first performed uniform quality control on each individual tumor from every dataset, filtering for primary IDH1^WT^ adult tumors and only neoplastic cells (Fig. 1a-c, SFig. 1a-d, STable1-3, Methods). All tumor cells were labeled as neoplastic by the original publications and confirmed to be malignant through in house inferCNV methods, which demonstrated canonical chromosome 7 amplification and chromosome 10 deletion (SFig. 1d, Methods). Cell types were annotated with adult and neurodevelopmental-like cell types, including those related to human cortical development such as radial glia (RG) and outer radial glia (oRG). Many of the cell types that we annotate in our meta-atlas have been described in GBM, such as mesenchymal (MES, expressing *CH13L1)*, astrocyte (expressing *GFAP* and *AQP4)*, immune reactive (*ANXA2, CD163),* oligodendrocyte (*OLIG2*), OPC (*PDGFRA*), neuronal populations (*MAP2, ESCAM, MIAT),* and dividing cells *(TOP2A* and *AURKB)*. We found multiple populations to express marker profiles reminiscent of neurodevelopmental cell types. These include Radial Glia (RG, expressing *ANXA2* and HES1)^14^, outer radial glia (oRG, expressing *HOPX* and *EGFR*)^4,21^, and the tri-IPC^22^ (*EGFR*, *PDGFRA*). Additionally, we observed the existence of a mesenchymal progenitor (MES.Progenitor, marked by *CHI13L1* and *SPON2*), an undefined progenitor, which exhibits characteristics of radial glia and a number of other mixed programs, and the NVP, which expresses radial glia markers such as *HOPX, HES1, GFAP, ANXA2* in addition to canonically perivascular markers such as *PDGFRB* and *NOTCH3* (Fig. 1b-c, STable2).

**Figure 1.**
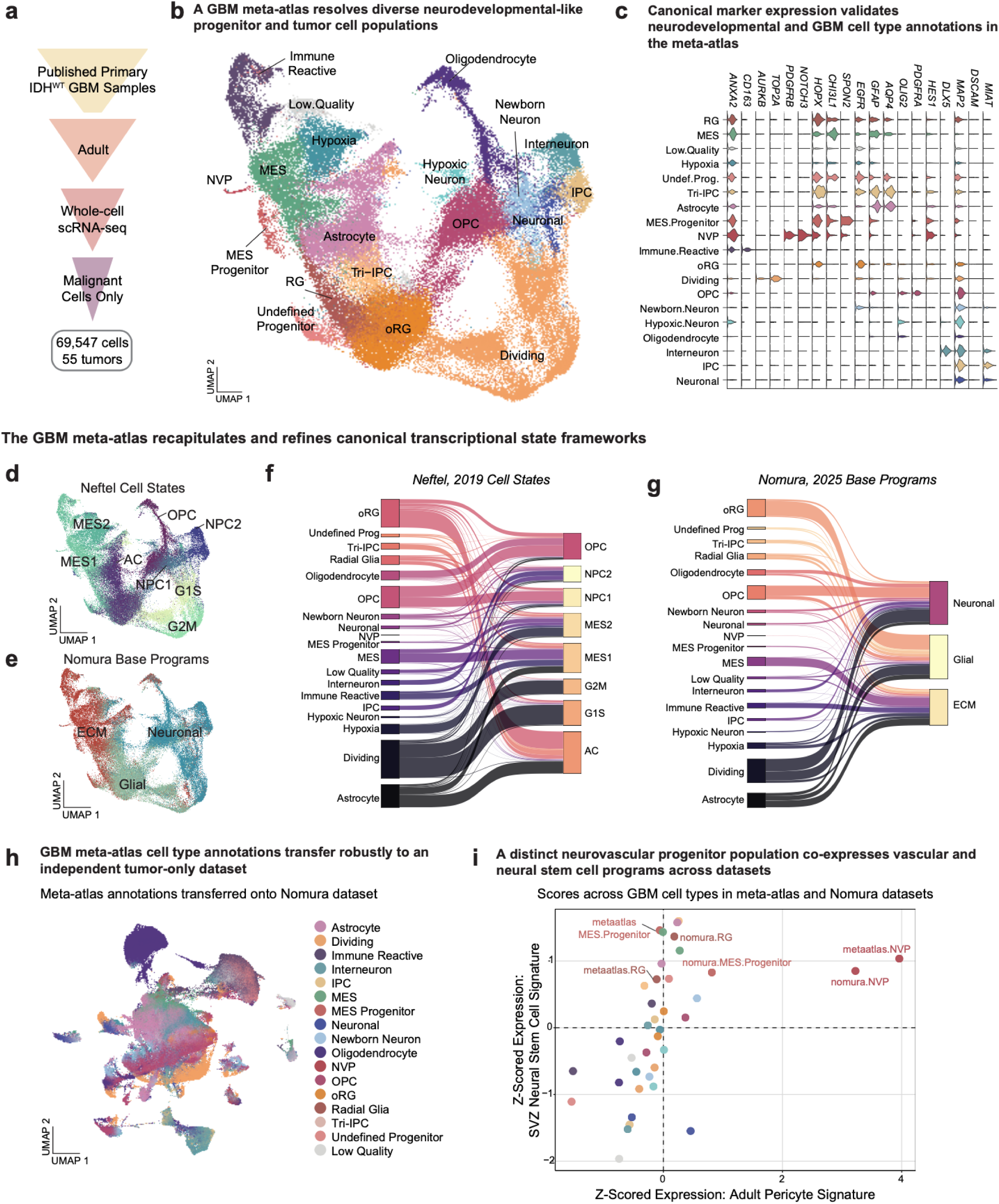
A GBM meta-atlas defines neurodevelopmental-like tumor progenitors and reveals a neurovascular progenitor with dual transcriptional identity. a) A stepwise exclusion process was implemented to curate the GBM meta-atlas from published primary GBM. The initial dataset included published primary GBM samples. The first criterion excluded pediatric samples, retaining only data from adult patients. Subsequently, samples harboring mutations in the isocitrate dehydrogenase 1 (IDH1) gene were excluded, focusing solely on IDH1 wild-type (IDH1^WT^) samples. The final exclusion step involved the removal of non-tumor cells per the original authors’ annotations. This filtration resulted in a final dataset comprising 69,547 cells across 55 tumors. b) Integrated Uniform Manifold Approximation and Projection (UMAP) visualization of malignant cells into a unified meta-atlas following stepwise exclusion criteria shown in panel a. Cells are colored by transcriptional cell type annotation derived from integrated clustering and marker-based annotation informed by adult and neurodevelopmental reference datasets (Methods). The meta-atlas resolves a spectrum of tumor populations, including differentiated or state-defined populations such as Mesenchymal (MES), Immune Reactive, Hypoxia, Dividing, Oligodendrocyte-like, Neuronal, Newborn Neuron, and Interneuron, as well as multiple progenitor populations with neurodevelopmental features. These include Radial Glia (RG), outer Radial Glia (oRG), Intermediate Progenitor Cells (IPC), Tri-lineage IPC (Tri-IPC), Mesenchymal Progenitors (MES.Progenitor), an Undefined Progenitor population, and a distinct Neurovascular Progenitor (NVP) population exhibiting mixed progenitor and perivascular transcriptional features. c) Expression of canonical marker genes across transcriptionally annotated glioblastoma (GBM) cell populations in the meta-atlas. Rows correspond to annotated cell types or states, and columns represent established markers of neurodevelopmental progenitors, GBM tumor states, and differentiated lineages. Radial glia (RG) and outer radial glia (oRG) populations express progenitor-associated markers including *ANXA2*, *HES1*, *HOPX*, *EGFR*, and GFAP, while intermediate progenitor populations (IPC and Tri-IPC) express *EOMES*-associated programs and cell cycle-linked genes. Mesenchymal populations (MES and MES.Progenitor) are enriched for CHI3L1 and SPON2, whereas astrocyte-like cells express *GFAP* and *AQP4*. The neurovascular progenitor (NVP) population uniquely co-expresses neural progenitor markers (for example, *HOPX*, *HES1*, *GFAP*) together with canonical perivascular markers including *PDGFRB* and *NOTCH3*. Cycling cells show elevated expression of proliferation markers such as *TOP2A* and *AURKB*, oligodendrocyte-lineage cells express *OLIG2* and *PDGFRA*, and neuronal populations express *MAP2*, *DSCAM*, and *MIAT*. d) UMAP visualization of GBM meta-atlas cells colored by cell state assignment based on Neftel 2019^10^ gene programs (Methods, STable3) e) UMAP visualization of GBM meta-atlas cells colored by cell state assignment based on Nomura 2025^20^ gene programs (Methods, STable3) f) Correspondence between meta-atlas cell type annotations (left) and cell state assignments (right). Band width represents the proportion of cells shared between each annotation pair in Neftel Cell States g) Correspondence between meta-atlas cell type annotations (left) and cell state assignments (right). Band width represents the proportion of cells shared between each annotation pair in Neftel Nomura Base Programs h) UMAP visualization of an independent tumor-only GBM dataset, calculated in house from the Nomura et al. data, colored by cell type labels transferred from the GBM meta-atlas. Meta-atlas annotations were projected onto the Nomura UMAP embedding using reference-based mapping. i) Scatter plot showing z-scored module activity (Methods) for an Adult pericyte module^23^ and an SVZ neural stem cell module^24^ across annotated GBM cell types in both the meta-atlas and Nomura datasets. NVP populations from both datasets exhibit high co-enrichment for the Adult pericyte and SVZ neural stem cell modules relative to other tumor cell types.

In order to benchmark our cell designations to previously published data, we leveraged existing sequencing efforts of GBM samples that have been influential in the field^10,20^. In these efforts, the authors define fundamental cell states occupied by GBM cells. We observe high concordance to these cell states and base programs with additional granularity. For example, the Neftel 2019 cell states defined as MES1 and MES1 map to our mesenchymal cells, hypoxia, and immune reactive populations. The Neftel astrocyte labels our astrocyte population, and OPC labels our oligodendrocyte cluster (Fig. 1d,f, STable3-4). We then asked how our granular designations correspond to the fundamental base programs defined by Nomura, et al^20^. Nomura extracellular matrix (ECM) defines our more mesenchymal populations, Glial labels our outer radial glia, radial glial, and Tri-IPC, and neuronal labels our neuronal and some progenitor subtypes (Fig. 1e, g, STable3).

Seeing that our meta-atlas cell states correspond highly, but with more granularity, to previously defined ones, we sought to validate the presence of our cell types in the recently published dataset. Using publicly accessible data, we performed in-house analysis of their tumor-only cells by following their stated threshold with minor changes to parameter stringency (Method). Of note, the authors of the Nomura, et al. study eliminated any tumor cells that were not completely certain to be tumor cells with copy number variation, resulting in 241,708 cells of the original ∼700k passing quality control. This is advantageous for our analysis as it gives us full confidence that all populations validated in their dataset are indeed tumor populations and not infiltrating populations from the nonneoplastic brain. We then labeled the filtered data, containing 241,708 malignant cells across 120 tumors with our annotations (Fig. 1h,e, STable5). Genescore correlation analysis determined the cross-applicability of our annotations to the new dataset, and we found that our labels transferred as expected. Notably, the NVP cell population from our meta-atlas and the Nomura data was among the most specific label transfers (SFig. 1h).

As we observed NVP to exhibit transcriptional profiles of perivascular and neural progenitor cells, we looked at the co-expression of transcriptional programs to determine if the co-expression we were observing is unique to the NVP population or if it is shared across GBM progenitor cell types. We scored all cell populations in the meta-atlas and Nomura datasets with adult brain pericyte signature^23^ and a developmental neural stem cell signature^24^. NVP cells in both datasets uniquely co-expressed the adult pericyte signature along with similar levels of neural stem cell signatures to other progenitors in the datasets such as radial glia and MES progenitors (Fig. 1i)

### NVP marker co-localization is observed across datasets and drives functional entropy

Within the pericyte and progenitor signatures, we identified several marker genes that can be used to combinatorially identify NVP. Within our annotated NVP population, we observed the co-expression of marker genes related to perivascular (*PDGFRB, NOTCH3, LAMA4)*, and neural stem cells (*NES, HOPX*) (Fig. 2a-b) across the three datasets; while these marker genes are also expressed in other cell types, they are uniquely co-expressed in NVP. Vascular cells derived from gliomas have been previously described^25,26^, and these tumor-derived endothelial populations have been shown to support tumor survival, even after treatment^27^. Moreover, PDGFRβ-positive tumor cells have been extensively characterized and even targeted therapeutically^28–38^, though none of these drugs are currently used as standard of care for patients. Additionally, some combinations of PDGFRβ-positive pericytes, vascular cells, and stemness markers such as *SOX2*^39,40^ and *NES*^41,41,42^ have been described in healthy and cancer contexts. The NVP cell type that we describe appears distinct from these previously described populations. Additionally, the co-expression of endothelial, mural, and neural progenitor features that we observed (Fig. 2a-h, Fig. 3c-g, SFig. 2a-b, SFig. 3) has not been characterized in the literature, but the plausibility that this population may exist is supported by these prior observations.

**Figure 2.**
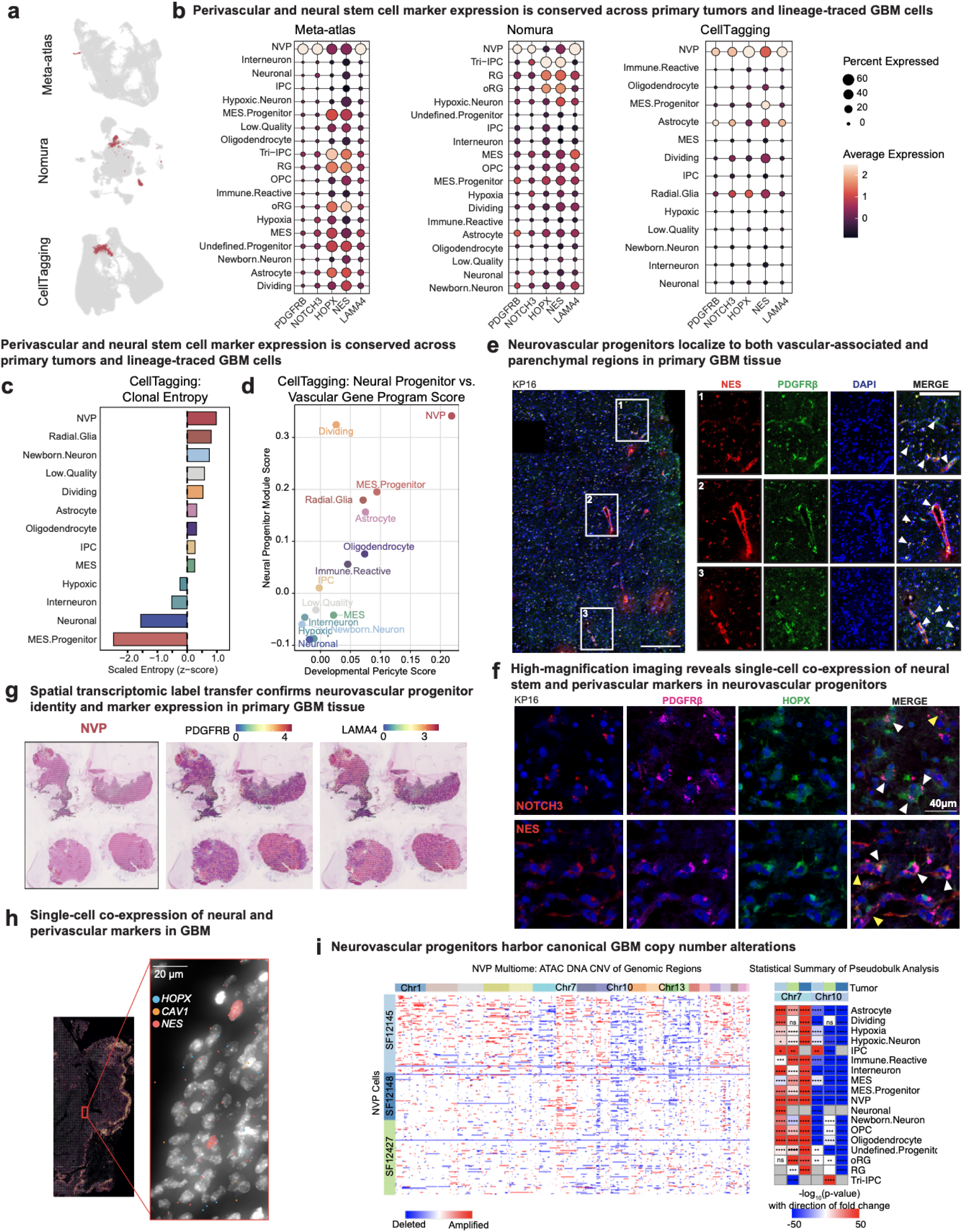
Lineage, spatial, and genomic analyses validate malignant neurovascular progenitors in GBM. a) UMAP visualizations of malignant GBM cells from the GBM meta-atlas, the independent tumor-only dataset from Nomura et al., and the CellTagging lineage-tracing dataset. Cells identified as neurovascular progenitors (NVP) based on meta-atlas-derived annotation criteria (Methods) are highlighted in red in each dataset. b) Cell type expression of perivascular markers (*PDGFRB*, *NOTCH3*, *LAMA4*) and neural progenitor markers (*HOPX*, *NES*) across annotated cell types in the GBM meta-atlas, the Nomura dataset, and the CellTagging^19^ dataset. While individual markers are detected across multiple tumor populations, their coordinated co-expression is most pronounced in the NVP population across all three datasets, supporting a conserved mixed transcriptional identity. c) Clonal entropy for each annotated tumor cell type in the CellTagging lineage-tracing dataset, displayed as scaled entropy (z-score). Clonal entropy quantifies the diversity of cell type partners co-occurring with a given cell type across inferred clones and was calculated from clone-level co-occurrence relationships using observed-versus-expected enrichment metrics (Methods). Higher entropy values indicate broader, less fate-restricted clonal connectivity. NVPs exhibit the highest clonal entropy among tumor cell types, consistent with extensive cross-lineage clonal relationships observed in lineage tracing. d) Z-scored module activity (Methods) for a developmental pericyte gene program^51^ and a neural progenitor gene program corresponding to dividing radial glia^67^ across annotated tumor cell types in the CellTagging dataset. Each point represents a transcriptionally defined cell type, positioned by its average module scores. NVPs exhibit coordinated enrichment for both developmental pericyte and neural progenitor programs relative to other tumor cell types, consistent with a mixed vascular-neural progenitor identity. e) *In situ* immunofluorescence was performed to assess co-localization of the perivascular marker PDGFRβ with the neural progenitor cell (NPC) marker NES in primary GBM tissue. Top row: 20× tile scan from tumor KP16 showing co-localization of NES (red) and PDGFRβ (green); nuclei are counterstained with DAPI (blue). Insets correspond to boxed regions in the overview image and display regions of marker co-expression, indicated by white arrows. Co-localization is observed both surrounding vascular structures and in tumor parenchyma. Scale bars: tile scans = 400 μm; inset panels = 200 μm (1.7X zoom). Representative images are shown. f) High-magnification immunofluorescence imaging demonstrates single-cell co-expression of perivascular and neural progenitor markers in neurovascular progenitors within primary GBM tissue. Representative 60× images show co-expression of NOTCH3 (red) and PDGFRβ (magenta) with the neural progenitor marker HOPX (green) (top row), and co-expression of PDGFRβ (magenta) with the neural progenitor marker NES (red) (bottom row). Nuclei are counterstained with DAPI (blue). White arrowheads indicate tumor cells exhibiting marker co-expression, whereas yellow arrowheads denote adjacent normal vascular structures. Scale bar = 40 μm. Images are representative of staining performed across three independent patient tumors. Representative images are shown. g) Representative Visium section^46^ (AT201_FO1_ii) annotated using RCTD-based label transfer from the GBM meta-atlas. Spots annotated as NVP are highlighted. Spatial feature plots for NVP markers *PDGFRB* and *LAMA4* are shown (Methods). h) Single-cell spatial co-expression of neural and perivascular markers in primary GBM (Region 05-0126-A2). Xenium-based spatial transcriptomic analysis showing transcript-level detection of *HOPX* (neural progenitor marker), *CAV1* (NVP-associated marker), and *NES* (neural stem cell marker). Left, overview of the Visium section with region of interest indicated. Right, high-magnification view demonstrating co-localization of neural and perivascular transcripts within individual nuclei. Outlined cells indicate representative triple-positive cells. Scale bar, 20 µm. i) Copy number variation (CNV) analysis was performed in joint scATAC-seq and scRNA-seq multiome profiling of 3 primary tumors. The scRNA data was used to identify cell types and the scATAC data was used to identify CNVs. Single-cell based analysis was performed with epiAneufinder^47^ and across all 3 tumors in the NVP, CNVs were observed including the characteristic chr 7 amplification (shown in heatmap on left in red) and chr 10 deletion (shown in blue). Pseudobulk analysis was also performed by combining the ATAC-sequencing from each cell type and calculating the deviation for each chromosome from diploid using a Mann-Whitney test. The -log10(p-value) for this calculation is shown, with the deleted chromosomes multiplied by negative 1 to yield a blue coloring in the heatmap on the right. Tumor colors match those from the heatmap^82^ on the left (SF12148: dark blue, SF12145: light blue, SF12427 green). Significance for each box indicates whether the amplification or deletion for that tumor and cell type is significant (ns = not significant, * < 0.05, ** < 0.0001, *** < 0.0000001, **** < 1e-10.

**Figure 3.**
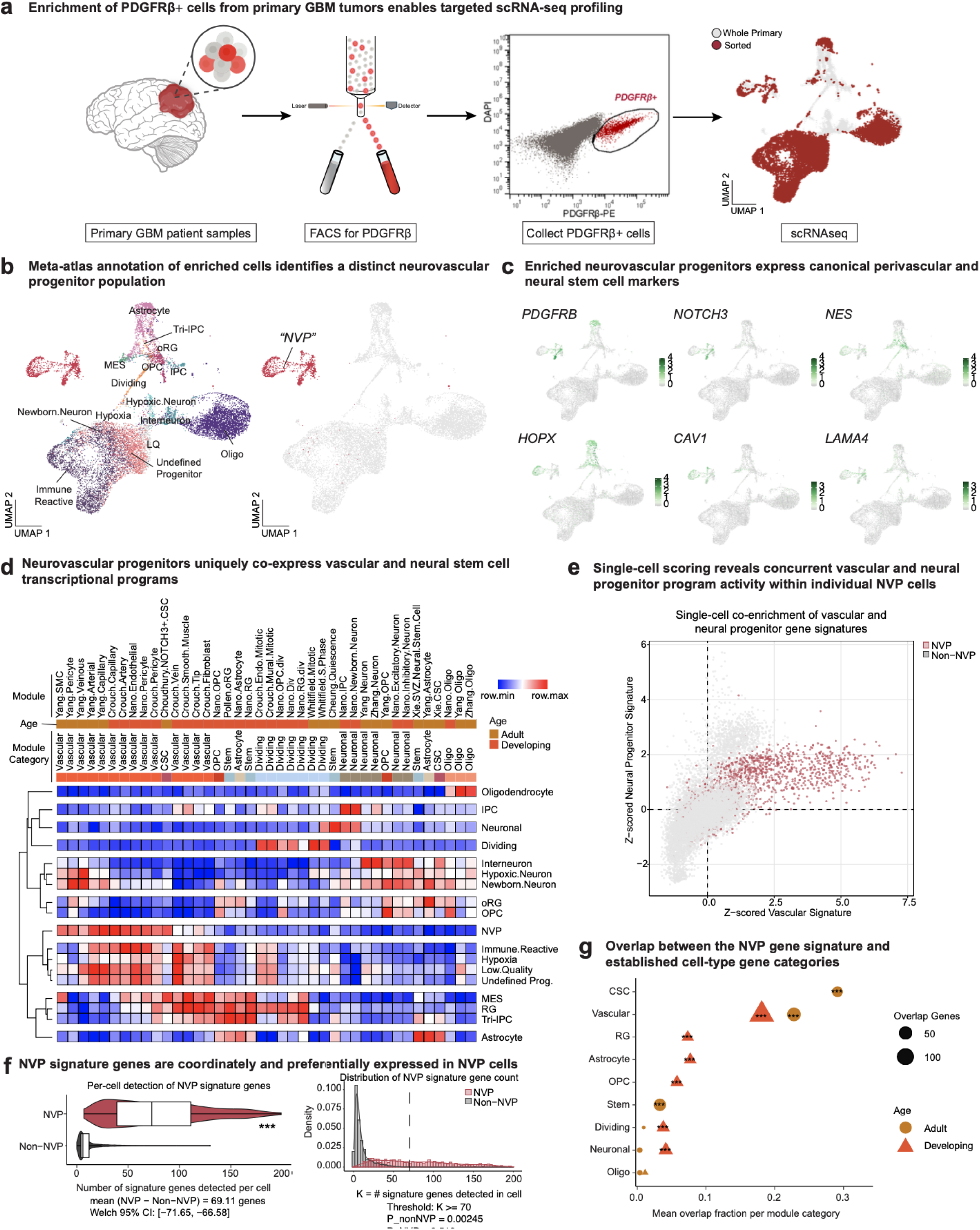
Enrichment of neurovascular progenitors from primary GBM tumors reveals a coherent mixed neurovascular transcriptional program. a) Schematic of fluorescence-activated cell sorting (FACS) and scRNA-seq enrichment strategy for primary GBM patient samples. Tumors (n=3) were dissociated into single-cell suspensions and stained with PDGFRβ-PE and DAPI to exclude dead cells. Live (DAPI-) PDGFRβ+ cells were isolated by FACS. FACS plot illustrates the gating strategy distinguishing PDGFRβ+ (red) from PDGFRβ-(gray) populations. Both sorted PDGFRβ+ cells and unsorted “Whole Primary” tumor cells were captured for scRNA-seq. UMAP shows the resulting distribution of Whole Primary (gray) and sorted PDGFRβ+ (red) cells following transcriptomic profiling (n = 15,996 cells). b) Left, Integrated UMAP of combined sorted and unsorted tumor cells annotated by projection onto the GBM meta-atlas (Methods), revealing diverse malignant cell populations. Right, cells annotated as neurovascular progenitors (NVP) are highlighted in red based on MapQuery label transfer from the GBM meta-atlas together with enrichment for vascular and neural progenitor gene programs (see panel c). This population forms a distinct cluster within the enriched dataset and corresponds to the putative NVP cell type. c) Feature plots showing expression of representative NVP and perivascular markers (*PDGFRB*, *NOTCH3*, *CAV1*, *LAMA4*) and neural progenitor markers (*NES*, *HOPX*) across the enriched scRNA-seq dataset. Marker expression is visualized on the UMAP embedding shown in panel b. These markers are co-expressed within the NVP cluster, whereas expression in other tumor cell populations is limited or non-overlapping, consistent with a mixed neurovascular transcriptional identity. d) Heatmap^82^ showing module scores for published vascular, neural progenitor, stem cell, and cell cycle gene programs across annotated tumor cell types from the enriched dataset shown in panel b. Cells were scored for each gene module using curated gene sets from published studies (Methods, STable4), and module scores were aggregated by cell type. Modules are grouped by developmental age (adult versus developing) and functional category (vascular, neural, stem, dividing, or lineage-specific), as indicated by the annotation bars.NVP exhibit concurrent enrichment of adult and developmental vascular programs alongside neural progenitor and stem cell programs, a pattern not observed in other tumor cell populations. e) Scatter plot showing z-scored module activity (Methods) for a developmental pericyte gene program (Crouch et al.) and a neural progenitor gene program corresponding to dividing radial glia (Nano et al.) in the PDGFRβ-enriched scRNA-seq dataset. Each point represents a single cell. NVPs (red, 3b) exhibit concurrent enrichment of both vascular and neural progenitor programs within individual cells, whereas non-NVP tumor cells (gray) generally lack coordinated co-expression of these signatures. Dashed lines indicate zero-centered z-scores for each module. f) NVP signature genes are coordinately and preferentially expressed in NVP cells. Left: Violin plots showing the distribution of the number of NVP signature genes detected per cell (defined as raw RNA counts > 0) in NVP and non-NVP cells. White boxes indicate the median and interquartile range. NVP cells exhibit a substantially higher number of detected signature genes compared to non-NVP cells, with a mean difference of 69.11 genes (Welch 95% confidence interval: [66.58, 71.65]). This large and tightly bounded separation motivated the use of K ≥ 70 detected genes as a conservative threshold for defining extreme co-expression in subsequent analyses (right). Right: Distribution of the number of NVP signature genes detected per cell (K) for NVP (red) and non-NVP (gray) populations. Dashed line indicates the threshold of K ≥ 70 detected signature genes. Empirical tail probability analysis shows that detection of ≥70 signature genes is rare in non-NVP cells (PDₒD₋□ᵥD = 0.00245), but frequent in NVP cells (PDᵥD = 0.519), demonstrating that high-level co-expression of the NVP gene signature is strongly enriched in NVP cells and unlikely to arise by chance in other tumor populations. g) Dot plot summarizing the overlap between the top 200 NVP signature genes and module-derived gene sets, stratified by functional category (Figure_Group) and developmental age (Figure_Age) as defined in (d). Each point represents one Figure_Group × Figure_Age category. The x-axis shows the mean fraction of genes within each category that overlap with the NVP signature, calculated after restricting all genes to the Seurat RNA feature universe (N = 30,670). Dot size reflects the number of overlapping NVP signature genes, and point shape denotes developmental age (adult or developing). Statistical enrichment was assessed using a one-sided hypergeometric test with the full RNA feature set as background. Significance is indicated as p < 0.05, p < 0.01, p < 0.001. Developing vascular and progenitor-associated categories show strong enrichment for the NVP signature, whereas adult oligodendrocyte-associated categories do not.

In addition to transcriptomic mixed identity, functional data from the CellTagging lineage tracing showed that NVP had the highest clonal entropy of any tumor cell type (Fig. 2c), exhibiting the highest diversity of clone partners. NVP from the CellTagging dataset similarly co-express neural progenitor and perivascular transcriptional programs (Fig. 2d). These data provided strong transcriptional and functional evidence of NVP significance, and we sought to further validate the distribution of NVP in primary tumor tissue. To examine the *in situ* distribution of NVP, we performed immunostaining for PDGFRβ, NOTCH3, with NES (common glioma stem cell marker^41^) and HOPX (marker for oRG^21^). We hypothesized that NVP cells might be spatially arranged near canonical perivascular populations; we thus explored co-localization of these marker genes in primary GBM tissue across tissue niches (Fig. 2e-f, SFig. 2b, SFig 3a-b). PDGFRβ and NES double-positive cells have been previously described in the context of tissue regeneration after injury^43,44^ and have been identified to be a characteristic of pericytes that can be reprogrammed into a neural lineage^45^, however this identity has not been described in the context of GBM. Because PDGFRβ has been well studied in GBM literature, we investigated another NVP marker, COL1A1, to examine if it had a similar co-localization profile across tumor sections with NES (SFig. 2a). Across these tumor two staining panels, we found that a subset of cells expressing pericyte markers exhibited co-expression with NOTCH3 and HOPX (SFig. 2b). Indeed, we observed that putative perivascular and NVP cells were seen in anatomical clusters or apart from other putative pericytes and in vessel-associated and parenchymal regions. We repeated this staining across 3 patient tumor samples and noted similar morphologies in samples from each patient.

Published spatial transcriptomic data^46^ was mapped to our GBM meta-atlas (Methods), enabling identification of NVP-annotated spots across multiple tumors (n = 3). NVP-labeled regions spatially coincided with expression of canonical NVP markers, demonstrating that the mixed neural–perivascular transcriptional program is spatially coherent within intact tumors (Fig. 2g). To resolve co-expression at true single-cell resolution, we analyzed Xenium multiplexed spatial transcriptomics. Across multiple tumors (n = 2), individual cells simultaneously expressed the NVP-associated marker CAV1 together with neural progenitor markers *HOPX* and *NES* (Fig. 2h). Detection of triple-positive cells confirms that neural and perivascular transcripts are co-expressed within the same malignant cells rather than reflecting spatial admixture of adjacent populations, providing orthogonal validation of the NVP state *in situ*.

Together, these data provided evidence that the co-expression of vascular and neural progenitor programs can be detected at the transcript and protein levels. The existence of NVP in not only our meta-atlas, but also the Nomura et al dataset and the CellTagging study provides strong evidence that NVP is indeed a neoplastic celltype. Therefore, to explore if NVP contains canonical GBM copy number variation, we sought to link the transcriptomic identity of the NVP cells to DNA aneuploidy within primary human GBM tumors by performing single-cell multiome sequencing on 3 primary tumor samples. First, we leveraged the single-cell transcriptomic data from this assay to annotate cell type based upon our meta-atlas, and across tumors we observed a subset of cells that were annotated as NVP. Next, we used epiAneufinder, a recently published tool to analyze CNVs from single-cell ATAC sequencing data^47^. Across all 3 tumors, it was able to identify chr7 amplification and chr10 deletions (Fig. 2i, SFig. 4) which are the most common in GBM^48^. When clustering by cell type, all of the tumor cell types intermixed across samples, suggesting that the CNVs were represented across cell types, including in NVP cells,, consistent with lineage relationships between cell types (SFig 4).

### FACS enrichment from primary tumors allows for molecular characterization of NVP

Given the presence of NVP cells as a small fraction of the tumor population, we sought to more extensively characterize this population. To do so, we employed a sorting strategy that would allow us to enrich for NVP. Direct-from-patient GBM samples were sorted for PDGFRβ expression immediately after surgical resection (without expansion or plating) using fluorescence activated cell sorting (FACS) (Fig 3a, SFig. 5a, STable6). PDGFRβ was selected based on our transcriptomic data as a high-sensitivity surface marker for NVP in order to provide the most enrichment possible (Fig. 1b-c, Fig. 2b, STable2). We therefore expected that our sorted dataset would contain other cell types. The sorted and unsorted cells acquired from (n = 3) patient tumor specimens were immediately captured for single-cell RNA-sequencing (scRNA-seq) (Fig 3a-c, SFig 5a-b). The resulting dataset consisted of 15,996 cells after quality control, including removal of doublets and copy number variation filtration for tumor cells (Methods, STable8). Tumor cells were annotated using the GBM meta-atlas, allowing us to molecularly define a subset of our sequenced cells as NVP (Fig. 3b-c, SFig. 5b). Notably, NVP cells do not express CD45 (*PTPRC*) or *SOX2* in the direct-from-patient context (SFig. 5c).

### Interrogation of vascular and neural stem cell gene program co-expression

We aggregated gene sets from published studies^23,24,49–52^ (STable4) to explore their activity in NVP cells and the rest of our data. NVP cells showed enrichment for adult vascular gene programs^23^, developmental vascular gene programs^51^, murine cancer stem cel^49^, normal stem cell programs^24,52^, and a recently identified meningioma NOTCH3+ stemness signature^50^ (Fig 3d). Vascular and neural progenitor signatures co-occur in the same cell in the NVP cluster but not other cell types (Fig. 3e). We generated a gene signature from the top 200 differentially expressed genes of the NVP cluster (STable7). Even with the expected dropout from single-cell analysis, NVP cells exhibit a substantially higher number of detected signature genes compared to non-NVP cells, with a mean difference of 69.11 genes (Welch 95% confidence interval: [66.58, 71.65]) (Fig. 1f, SFig. 5d). This analysis re-emphasizes in our sorted cell population that NVP marker genes are uniquely co-expressed within NVP cells, defining this malignant population. The NVP signature exhibits the most robust overlap with published cancer stem cell (CSC) signatures in addition to adult and developing vascular and developing radial glia gene programs (Fig. 3g). We tested the ability of NVP cells to alone form vascular structures and observed they do not show evidence of tube formation capability (SFig. 5e-f), leading us to further focus on their putative function as a cancer stem-like population.

### DNA barcoding-based clonal analysis of enriched NVP cells from human GBM samples demonstrates dual fate of NVP

Given strong evidence that NVP exhibits progenitor cell behavior, we sought to establish where NVP fits into the previously-established hierarchy of GBM lineages. Studying human tumors involves additional challenges compared to mouse *in vivo* systems, including the observation that *in vitro* expansion of primary tumors alters tumor composition and may result in decreased progenitor complexity^53^. To address these challenges, we recently developed and optimized a human organoid tumor transplantation (HOTT) system that both preserves tumor heterogeneity and transcriptional fidelity to the parent tumor while also modeling key features of the tumor microenvironment^54^. Briefly, tumor cells directly from the patient are labeled with lentivirus containing GFP and transplanted into an already existing cortical organoid that models key features of the developing human cortex. A unique feature of this HOTT system is that it can also enable the exploration of isolated subsets of tumor cells and allows for additional molecular modifications to be introduced. We took advantage of these features by repeating our enrichment for NVP by sorting for PDGFRβ+ cells and then performed clonal tracking using the well-described CellTag DNA barcoding system^55,56^ (Fig. 4a). In parallel, we validated that the PDGFRβ-fraction was depleted of NVP cells (SFig. 7a-b).

**Figure 4.**
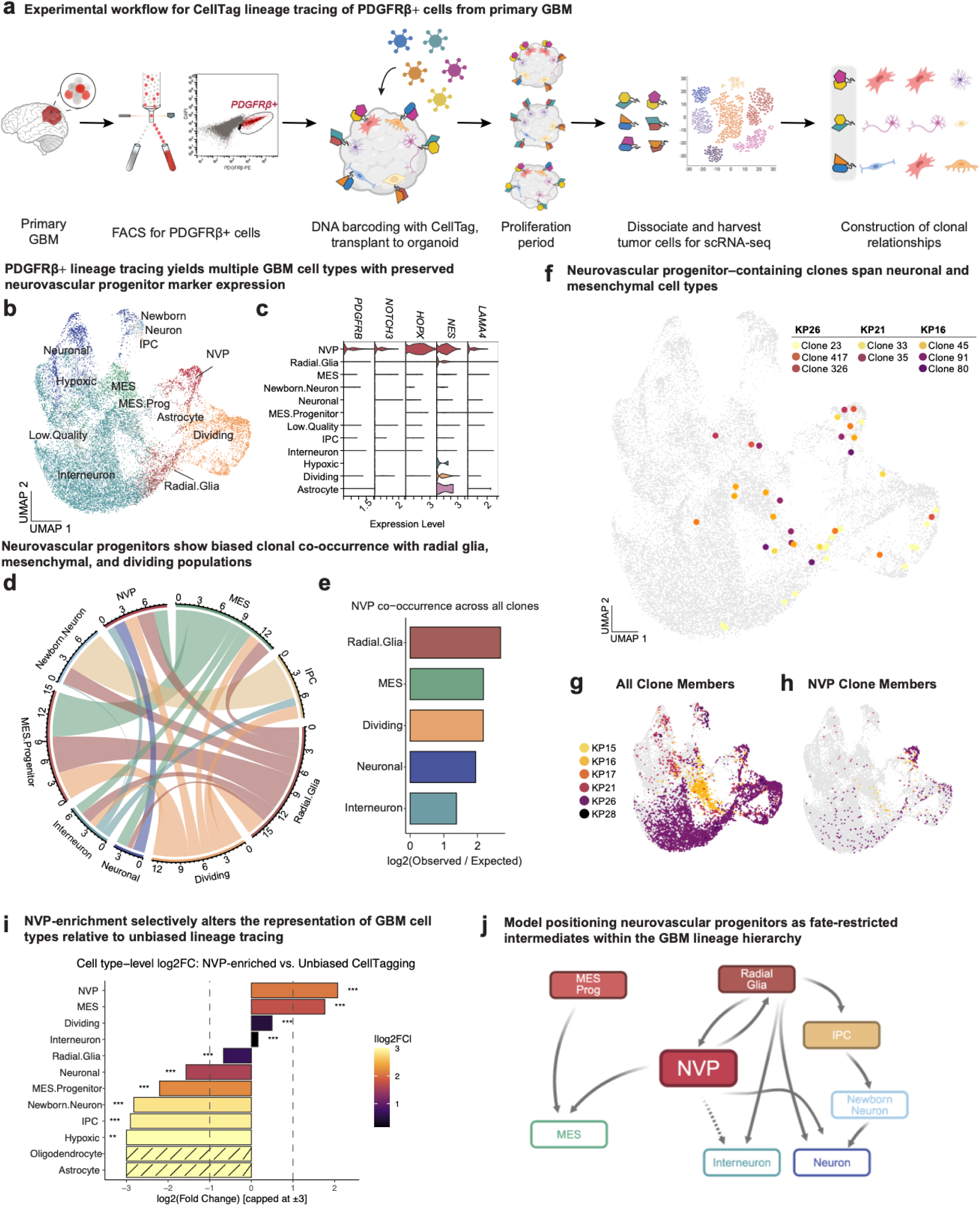
Lineage tracing of PDGFRβ+ neurovascular progenitors reveals restricted neuronal and mesenchymal clonal partnerships in GBM. a) Primary GBM samples (n = 6) were subjected to FACS isolation of DAPI-PDGFRβ+ cells. These cells were then transduced with a lentiviral library containing plasmids which contain unique CellTag DNA barcodes and GFP. After transduction, cells were transplanted onto week 8-12 human cortical organoids. Following a proliferation period of 12-18 days, the tumor cells were dissociated from the organoid and sorted based on GFP. GFP+ cells are harvested for scRNA-seq. The resultant data include paired CellTag DNA barcode information with transcriptomic profiles, allowing the construction of clonal relationships (Methods). b) UMAP visualization of PDGFRβ⁺-enriched tumor cells (n = 20,153 cells) following transplantation into human cortical organoids (HOTT). Cell type identities were assigned by projecting single-cell transcriptomes onto the reference CellTagging^19^ dataset using MapQuery (Methods). The enriched population resolves into multiple canonical GBM tumor cell states, including neuronal, mesenchymal (MES), radial glia-like, and dividing populations. NVP cells form a distinct cluster within the enriched dataset and retain co-expression of NVPr markers. c) Expression of key perivascular (PDGFRB, NOTCH3, LAMA4) and neural progenitor (HOPX, NES) markers across annotated cell types, confirming preservation of the NVP transcriptional program in PDGFRβ⁺-enriched cells. d) Chord diagram summarizing enriched clone-level co-occurrence relationships among transcriptionally defined GBM cell types across all CellTag-traced clones (Methods). Ribbon width reflects enrichment magnitude, and sectors are colored by cell type. Low.Quality annotations were excluded. NVP cells show preferential clonal associations with radial glia-like, mesenchymal (MES), and dividing populations. e) Log₂ enrichment of clone-level co-occurrence between NVP cells and other GBM cell types (Methods). Only cell types observed in at least 10 NVP-containing clones are shown. Positive values indicate preferential co-occurrence with NVP within clones. f) Representative CellTag-defined clones that contain at least one NVP cell. Individual clones are overlaid on the global transcriptional embedding, with clone members colored by patient of origin. Selected clones from three tumors (KP26, KP21, KP16). g) All clone members (left) colored by patient. h) Clones containing at least at least 1 NVP cell colored by patient. i) Horizontal bars show the log₂ fold change in relative abundance for each cell type, computed as log₂(Freq_{NVP-enriched Dataset} / Freq_{Unbiased CellTagging Dataset}) and capped at ±3 for visualization. Color intensity reflects the absolute magnitude of the effect size (|log₂FC|) independent of direction. Dashed reference lines at +1 and −1 indicate two-fold enrichment or depletion, respectively. Asterisks indicate significance from per-cell type two-sample proportion tests with Benjamini-Hochberg FDR correction across cell types (* FDR < 0.05; ** FDR < 0.01; *** FDR < 0.001). No asterisks are shown for cell types that are completely absent in one dataset. Hashed (striped) bars indicate structural absence (present in the Unbiased CellTagging Dataset but not detected in the NVP-enriched Dataset). j) Model positioning NVP as fate-restricted intermediates within the GBM lineage hierarchy. Schematic summarizing inferred relationships from PDGFRβ⁺-enriched CellTag lineage tracing and clonal co-occurrence analyses (Fig. 4C-I; Methods). NVP is depicted as an intermediate state with restricted fate potential, exhibiting clonal connections to both neuronal lineages (radial glia/IPC leading to newborn neurons and neurons) and mesenchymal (MES) programs. Arrows indicate favored clone partnerships and putative differentiation trajectories observed in the HOTT CellTag experiments, rather than definitive developmental directionality.

Upon transplantation of the NVP-enriched tumor fraction into HOTT, we observed the recapitulation of complex cell morphology that we had previously seen *in situ* (SFig. 6a). To perform this experiment, we first infected freshly dissociated GBM cells from 6 patients with a lentiviral library containing scRNAseq compatible CellTag barcodes and a GFP tag before transplanting onto cortical organoids. We delivered the virus at a multiplicity of infection that would result in 3 - 4 barcodes introduced in each cell, enabling high-confidence clone calling in subsequent analysis (Methods). We then allowed the tumor cells to proliferate before harvesting for scRNA-seq. The paired analysis of CellTag barcodes and transcriptional signatures provides the identity of tumor cells derived from the same clone.

As before, we performed quality control analysis, doublet removal and filtered for tumor cells using copy number variation analysis, yielding 20,153 cells across 6 unique tumors. Cells were uniformly labeled with CellTag-containing lentivirus (SFig. 6b), giving all cells the opportunity to be included in a clone. Our analysis showed that across our 6 tumors, we averaged a multiplicity of infection of 2-3, meaning that the probability of any two cells randomly receiving the same combination of barcodes is infinitesimally small, as has been previously described^56^. Using the CellTag clone calling pipeline^55^, we identified 1465 clones ranging in membership from 2 to 28 cells, with an average size around 3 (SFig. 6c, STable9-10). The transcriptional analysis was performed orthogonal to the clonal analysis, and identified a subset of cell types that we typically observe within GBM tumors (Fig. 4b-c, SFig. 6d-e, STable10). NVP cells in this data reflect marker expression profiles consistent with the GBM meta-atlas, Nomura, and the unenriched CellTagging dataset (Fig. 4c, SFig. 6f-g).

After performing clonal analysis on each tumor individually, we merged all tumors and clone identities in order to visualize trends in clone membership on the same UMAP space. Each clone represents a set of cells that are derived from a common progenitor in the HOTT system, though the parent cell may or may not remain or be captured by our system. As such, we present the clones from our analysis in sum (Fig. 4g) and filtered for those that are restricted to containing one or more NVP cells in the clone (Fig. 4h) bottom right, UCSC Genome Browser: https://gbm-nvp.cells.ucsc.edu). While here we pictorially represent select representations of clones with interesting and unexpected cell type membership (Fig. 4f), the vast majority of the total clones were comprised of cells in the same cluster, especially that of cycling cells (STable9-10). Because having multiple clone members of the same cell type is more likely than having clone members of different cell types, this provides confidence that the clone calling strategy indeed resulted in related cells being highlighted. Additionally, multiple cell types such as the “Immune Reactive” population did not participate at all in clones at all, adding confidence to the specificity of clone calling.

Upon analyzing the cell types represented in clones containing NVP cells, we observed that Radial Glia and MES cells as the top clone partners (Fig. 4d,e,h). Our informatic analysis and *in situ* validation led us to hypothesize that NVP cells have the propensity to generate both neuronal and MES cells within the tumor. Indeed, we found several clones across multiple tumors that contained an NVP cell and at least one neuronal and/or MES cell, confirming the dual potency of this population. Because of the potential for nonspecific sorting with our enrichment strategy, only clones that include an NVP cell were considered to be clonally related to NVP. Notably, this eliminates our ability to detect symmetric divisions that no longer retain an NVP progenitor in the clone (Fig 4h). While we previously demonstrated GBM cell type composition changes as a result of NVP depletion, this direct-from-patient clone interrogation advances our understanding of NVP function by providing a concrete relationship between multiple tumor cell types. We note that across canonical models of GBM, neuronal and MES populations are thought to sit at opposite poles of GBM transcriptional identities^12,57^, highlighting a unique aspect of NVP cells in that they are capable of giving rise to divergent states within the tumor (Fig. 4j).

### NVP-enriched samples can generate *a majority* of the parent tumor

To assess how progenitor-intrinsic potential versus tumor context shapes lineage output, we directly compared the cellular composition recovered from PDGFRβ+ NVP-enriched lineage tracing to that obtained from our parallel, unbiased CellTagging study, focusing on the three tumors profiled in both experiments. This analysis revealed that enrichment for NVP substantially and selectively altered the spectrum of tumor cell types generated, rather than simply amplifying a scaled-down version of the parent tumor. While NVP-enriched cultures were, as expected, significantly enriched for NVP itself and for specific progenitor-adjacent and lineage-related populations, multiple tumor cell types readily detected in the unbiased lineage tracing experiment were markedly depleted or entirely absent (Fig. 4i). These shifts indicate that although NVP possesses an intrinsic capacity to function as a progenitor and generate both neuronal and mesenchymal lineages, the full realization of tumor cell diversity depends strongly on interactions with other tumor cell types present at initiation. Together, these results position NVP as a fate-restricted but highly influential intermediate within the GBM hierarchy whose lineage output should be understood within the context of the broader tumor cell population.

### NVP-depletion remodels tumor state composition and reveals NVP-centered signaling networks

To directly test how depletion of the PDGFRβ+ fraction reshapes tumor composition and to determine whether NVP-associated programs influence other malignant states, we performed parallel lineage tracing on matched CD45-PDGFRβ+ (POS) and CD45-PDGFRβ-(NEG) fractions isolated by FACS from (n = 3) primary IDH1^WT^ GBM tumors (SFig. 7a,c). As expected, the POS input was strongly enriched for transcriptionally defined NVP cells relative to unsorted tumor, whereas the NEG fraction was depleted of NVP at baseline (SFig. 7b). Each fraction was independently CellTagged and transplanted into the HOTT system prior to scRNA-seq (n = 22,321 cells), enabling direct comparison of lineage output and state composition between PDGFRβ-enriched and PDGFRβ-depleted initiators (SFig. 7c–f, STable11-12).

Across tumors, POS- and NEG-derived cells segregated by condition while retaining comparable patient representation (SFig. 7d,e). Analysis restricted to clone-member cells revealed marked shifts in cell-type representation following PDGFRβ depletion (SFig. 7f-g). Compared to the depleted condition, the POS condition showed increased representation of Astrocytes, Radial.Glia, NVP, Dividing cells, Interneurons, MES, and Newborn Neurons. IPC and Neuronal cells were more prevalent in the NEG condition compared to the POS. Clone co-occurrence networks (n = 1592 clones) further demonstrated condition-specific remodeling: POS-enriched clones showed strong NVP-centered interactions, whereas NEG-enriched clones displayed increased co-occurrence among alternative progenitors, including MES.Progenitor, Radial Glia, and IPC populations (SFig. 7h-i). Notably, despite initial depletion, NVP-like cells re-emerged within NEG-derived cultures, indicating that NVP-associated transcriptional identity can arise from the cell types present in the NEG condition.

Given these compositional and clonal shifts, we next examined the function of NVP in mediating tumor composition. Ligand–receptor modeling across all lineage-traced cells positioned NVP as a central signaling node characterized by high outgoing and substantial incoming interaction and strength (SFig. 7j). NVP demonstrated broad engagement in contact-dependent receptor–ligand interactions, including adhesion-associated modules (e.g., JAM, NGL, CADM families) and NOTCH signaling (SFig. 7k-l). Pathway-level analyses highlighted druggable contact-mediated programs such as ADGRB, CD46, GAP, NOTCH, NGL, and JAM in which NVP acted as a major signaling source to multiple recipient states.

Together, these findings indicate that depletion of the PDGFRβ+ compartment does not simply reduce downstream lineages but instead remodels progenitor balance and communication architecture. Beyond functioning as a lineage intermediate, NVP occupies a hub-like position within the tumor signaling network, coordinating contact-dependent interactions that shape GBM state composition.

### Identification and elimination of NVP in an *in vivo* developmental GBM model extends survival

Our data suggests a strong role for NVP in human tumors, and we wanted to validate whether NVP cells are functionally important *in vivo.* Given the link to developmental cell types and progenitors in GBM, we explored scRNA-seq data from a published, tractable, developmentally derived *in vivo* model of GBM^58,59^. Excitingly, this model of GBM allows exploration of diverse progenitor populations that also exist in human patient tumors. This approach uses *in utero* electroporation to knockdown three tumor suppressors using CRISPR-Cas9, giving rise to GBM-like tumors postnatally. We re-analyzed these data from the lens of our human-annotated GBM cell types, finding numerous populations of interest including a putative NVP that resembled the NVP population (Fig. 5a-b). To more formally assess correspondence between mouse tumor populations and human GBM cell states, we computed gene score correlations between mouse clusters and human meta-atlas-defined programs. This analysis demonstrated that the putative mouse NVP population showed the highest concordance with the human NVP program, supporting cross-species conservation of this cell state within GBM (Fig. 5b, Methods). Identifying NVP in this *in vivo* model of GBM provided orthogonal confirmation of the significance of NVP cells and provided a system in which to manipulate this population. Therefore, we performed *in utero* electroporation using this established *3xCr* approach which initiates tumors with CRISPR-mediated knockdown of *Pten*, *Nf1*, and *Trp53*.

**Figure 5.**
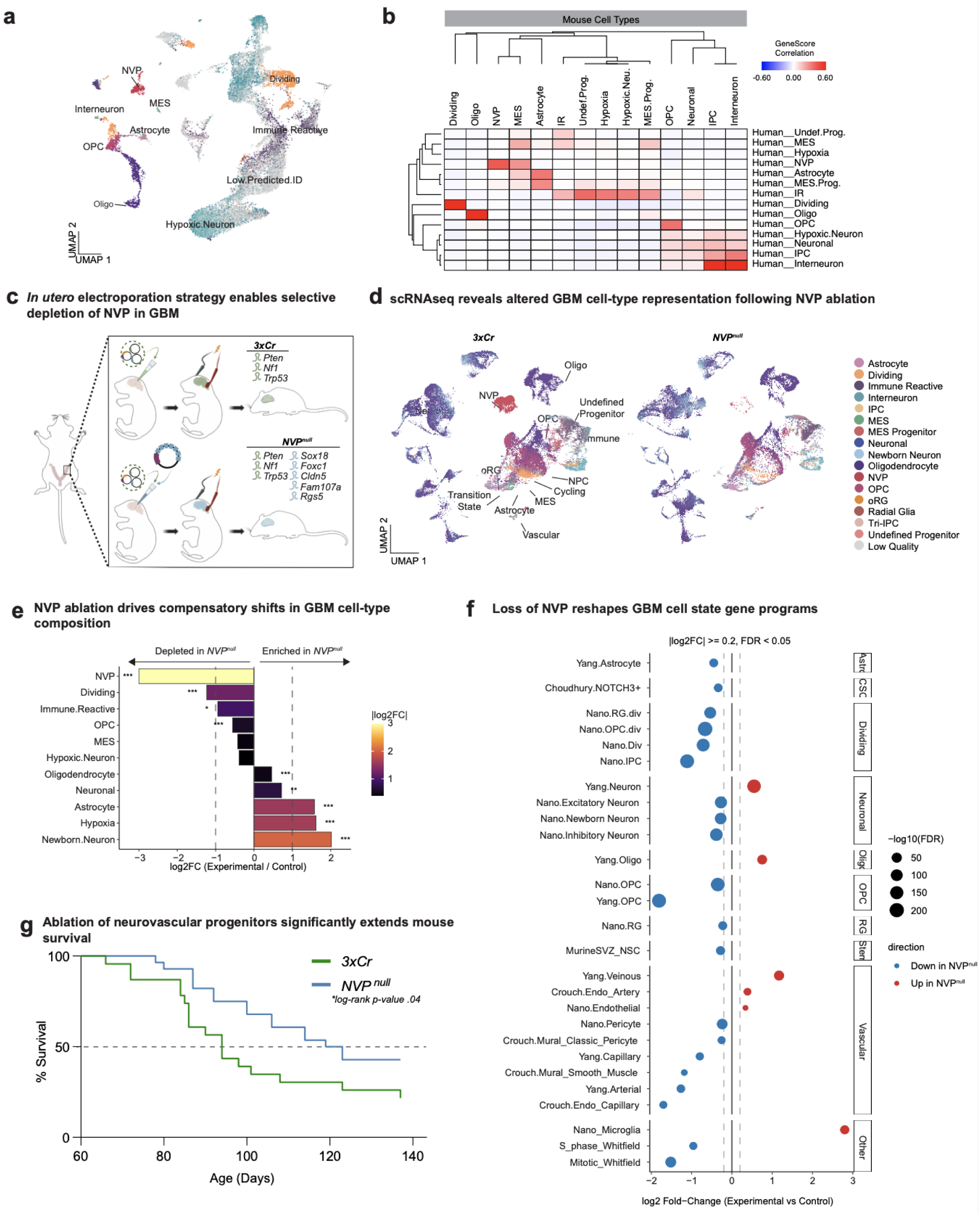
*In vivo* ablation of GBM neurovascular progenitors alters tumor composition and extends survival. a) scRNA-seq data from a developmental mouse GBM model generated by *in utero* electroporation^58^. Cells were processed using the standard analysis pipeline and annotated by projection onto the human GBM meta-atlas using MapQuery (Methods). Major tumor and neural cell populations are labeled, including a distinct cluster corresponding to NVPs. Cells with low annotation confidence following cross-species label transfer are indicated as Low Predicted ID. b) Heatmap^82^ showing gene-score correlations between transcriptionally defined mouse GBM cell populations (columns) and human GBM meta-atlas cell types (rows). Gene scores were calculated from cluster marker genes and used to compute pairwise Pearson correlations across shared genes (Methods). Hierarchical clustering was applied to both axes. The mouse cluster corresponding to the NVP population shows the strongest correlation with the human NVP signature, supporting the presence of an NVP-like cell state in this developmenta*l in vivo* GBM model. c) Schematic of the *in utero* electroporation (IUE)-based developmental mouse GBM model used to selectively deplete NVPs. Control tumors were generated by CRISPR-mediated knockout of Pten, *Nf1*, and *Trp53* (*3xCr*). For NVP depletion (*NVP^nul^*^l^), the same 3xCr background was combined with simultaneous targeting of five NVP-enriched genes (Sox18, Foxc1, Cldn5, Fam107a, and Rgs5), selected based on their specificity to the NVP cluster identified in panel a and minimal expression in other tumor cell types (Methods, SFig. 8) d) UMAP visualization of (n = 80,530) tumor cells from (n=4) control (*3xCr*) and (n=4) NVP-depleted (*NVP^nul^*) tumors profiled by single-nucleus RNA-sequencing. Cells are colored by transcriptional identity assigned via projection onto the GBM meta-atlas (Methods). In control tumors, a discrete NVP population is readily detected, whereas NVP cells are nearly absent following targeted depletion. e) Horizontal bars show the log₂ fold change in relative abundance for each tumor cell type in NVPⁿᵘˡˡ versus 3xCr control tumors, computed as log₂(experimental fraction / control fraction) and capped at ±3 for visualization (Methods). Bar color encodes the absolute effect size (|log₂FC|; inferno scale), independent of direction. Dashed reference lines at log₂FC = ±1 indicate two-fold depletion or enrichment. Asterisks denote significance from two-sample proportion tests with Benjamini-Hochberg FDR correction across cell types (* FDR < 0.05; ** FDR < 0.01; *** FDR < 0.001), with significance suppressed for categories structurally absent in one condition. Striped bars indicate cell types not detected in one condition. Cell types with fewer than 10 cells in either experimental or control tumors were excluded from the analysis. f) Differential activity of published gene programs after *in vivo* NVP depletion. Points show gene program module scores (Methods) comparing NVPⁿᵘˡˡ (Experimental) versus *3xCr* (Control) tumors, summarized as log₂ fold-change (Experimental/Control) across tumor cells (Methods). Only modules meeting |log₂FC| ≥ 0.2 and BH-FDR < 0.05 are shown; point size denotes significance (−log₁₀(FDR)) and color indicates direction (blue, decreased in NVPⁿᵘˡˡ; red, increased). Modules are grouped by functional class (right) g) Kaplan-Meier curve showing survival of *3xCr* (n=22) and *NVP^null^* (n=28) mice. The *NVP^null^* cohort showed an increase in median overall survival (50% survival depicted by dashed line) compared to the 3xCr cohort (116 vs. 94 days, log-rank p-value = .04).

Using this developmental *in vivo* tumor system, we sought to explore how NVP contributes to GBM composition by knocking NVP out *in vivo*. Although PDGFRβ is an excellent surface marker for enrichment of NVP cells, in both our meta-atlas and our sorted cell analysis we noted that it was not entirely specific to the NVP population. Thus, the genes for knockout were chosen based upon both their expression in the human NVP cells and their specificity to the mouse NVP cluster. Notably, based upon the RNA expression, these genes used to knockout NVP were not expressed in other cell types (SFig. 8a). These included highly specific transcription factors (*Sox18* and *Focx1* with known roles in endothelial differentiation^60,61^), neural stem cell marker gene *Fam107a*, endothelial marker *Cldn5* and mural marker *Rgs5* (SFig. 8a). Although many of these genes have known roles in vascular development, remodeling, or cancer, no coordinated functional role has been described characterizing their ability to initiate or maintain progenitor cell types in glioblastoma. Our experimental condition applied the same *3xCr* approach but added in simultaneous knockdown of our five NVP targets (*NVP^null^*), enabling exploration of how the resultant tumors would differ without NVP (Fig. 5c).

Our initial analysis of this model indicated robust and consistent representation of the NVP population at the postnatal day 70 (P70), thus we sacrificed 4 control *3xCr* and 4 experimental *NVP^null^* mice with equal numbers of male and female animals at this timepoint (P70) to explore the impact of NVP on tumor cell type composition (SFig 8b). We performed single-nuclei RNA-sequencing (snRNA-seq) and obtained data from 80,530 nuclei after quality control (Fig. 5, SFig. 8b, STable13). Specific target depletion was verified at the protein level (SFig. 8c-g) and the specificity of sgRNA activity was verified via whole exome sequencing (SFig. 8g). As before, we annotated cell types with our meta-atlas reference, observing that the *NVP^null^* resulted in a 96% reduction of the NVP cell fraction (Fig. 5d-e). Thus, we calculated the cell type composition for each of the 8 profiled tumors and compared how they differed with and without NVP to explore how NVP functionally contributes to GBM tumor biology.

Loss of NVP resulted in marked shifts in tumor cell-type composition. Cycling cells and OPCs were significantly depleted in the experimental condition. In contrast, cells from the neuronal lineage, hypoxic cells, astrocytes, and more differentiated oligodendrocytes increased relative to control (Fig. 5e). Consistent with the observed shifts in tumor cell-type composition, ablation of NVPs also resulted in widespread remodeling of cell-state-specific transcriptional programs. Gene program analysis revealed a coordinated reduction in vascular-associated and neuronal programs in *NVP^null^*tumors, including endothelial, mural, and neural lineage signatures that were prominent in control tumors. In parallel, alternative progenitor- and differentiation-associated programs were selectively enriched following NVP loss, indicating activation of compensatory cell states rather than uniform suppression of tumor identity (Fig. 5f). Notably, these transcriptional changes mirrored the compositional alterations observed at the single-cell level, linking NVP ablation to both loss of vascular-progenitor-associated gene programs and emergence of distinct neural and progenitor-like states.

Finally, we evaluated the survival of a separate cohort of 3*xCr* (n=23) and *NVP^null^*(n=28) mice. The *NVP^null^* cohort showed an increase in median overall survival compared to the *3xCr* cohort (119 vs. 91.5 days, *log-rank p-value = .03*) (Fig. 5e), indicating that the shifts in cycling populations and cell populations are sufficient to impact tumor growth and *in vivo* survival.

Together, these findings position NVPs as a conserved, functionally important, and fate-restricted intermediary within the GBM lineage hierarchy. While NVPs do not generate all tumor cell types, their selective ablation disrupts key proliferative and vascular programs, triggers compensatory differentiation, and significantly prolongs survival, highlighting their central role in shaping GBM tumor architecture.

## Discussion

A central challenge in GBM biology is determining how transcriptionally defined heterogeneity translates into functional tumor behavior. Although single-cell atlases have established that GBM contains multiple progenitor-like populations, they cannot resolve lineage directionality or whether progenitors can traverse distinct developmental programs. Here, we identify and functionally characterize a tumor-progenitor population that occupies an intermediate position within established GBM hierarchies and exhibits an exceptional capacity to cross lineage boundaries. We adopted a progenitor-centric strategy by constructing a GBM-specific meta-atlas restricted to adult, IDH1 wild-type, direct-from-patient tumors. This approach was motivated by prior work demonstrating that progenitor populations disproportionately shape tumor propagation and plasticity^4,6,10,57^. Integrating multiple datasets under uniform annotation ensured that identified populations represent conserved, tumor-intrinsic cell types rather than dataset-specific artifacts or microenvironmental contamination.

Within this framework, we identified the neurovascular progenitor (NVP), a malignant cell population that co-expresses neural progenitor and perivascular transcriptional programs. NVP cells were consistently detected across independent datasets, retained canonical GBM copy number alterations, and showed high specificity in label transfer to a stringently filtered tumor-only dataset ^20^, supporting their classification as a bona fide tumor cell type. Crucially, NVP is distinguished by its functional behavior. Lineage tracing in the companion CellTagging study revealed that progenitor populations differ in their ability to cross organizational axes of tumor hierarchy, with NVP exhibiting the highest degree of cross-lineage clonal connectivity. Direct clonal analysis in the present study demonstrated that individual NVP cells can generate both neural-like and mesenchymal tumor populations across patients. This dual-fate capacity cannot be inferred from transcriptional data alone and distinguishes NVP from other stem-like progenitors that remain largely fate-restricted.

These findings help reconcile prior models of GBM organization. Single-cell studies propose a tri-lineage hierarchy in which neuronal-proneural, oligodendroglial, and astrocytic-mesenchymal states coexist in dynamic equilibrium yet exhibit strong proneural-mesenchymal antagonism ^12^. Cell-of-origin studies further show that progenitors generate tumors with durable lineage signatures ^57,62^, raising the question of whether progenitors within established tumors retain multi-lineage potential. Our direct clonal links between NVP, radial glia-like cells, and mesenchymal populations identify NVP as a population capable of traversing this divide, providing a mechanism for maintaining antagonistic transcriptional states within the same tumor. Our results also contrast with lineage-of-origin frameworks proposing distinct perivascular malignant lineages. Previous work ^63^ has suggested that GBM aligns along radial glia-derived neural and neural crest-derived perivascular lineages. Rather than supporting a separate perivascular lineage, our data indicate that NVP is a tumor-intrinsic progenitor occupying an intermediate hierarchical position that clonally links proneural and mesenchymal compartments within established GBM hierarchies.

Spatial, molecular, and functional validation further support this interpretation. NVP cells localize to both vessel-associated and parenchymal regions and co-express perivascular and neural progenitor markers, yet they do not form vascular structures in isolation and lack features of canonical endothelial or mural cells. Their defining property is functional: the ability to generate diverse tumor cell types and act as a central lineage-intermediate population.

Despite this breadth, NVP is not sufficient to generate the entire tumor. Enrichment and depletion experiments show that NVP contributes substantially but incompletely to tumor composition, and that other progenitors can compensate for its loss. This is notably observed in our *in vivo* experiment, where ablation of NVP cells extends lifespan significantly, but the mice ultimately do succumb to the tumor. This distributed hierarchy mirrors principles of neurodevelopment, where intrinsic progenitor programs establish competence but extrinsic context refines fate. The re-emergence of NVP from depleted fractions and its role as a signaling hub further underscore the limitations of studying isolated cell types outside their native tumor ecosystem. Together, these findings position NVP at a functional nexus within GBM hierarchies: a progenitor with restricted yet unusually broad fate potential that links divergent lineage programs and contributes disproportionately to tumor plasticity. More broadly, this work illustrates how integrating lineage tracing with multi-dataset validation moves beyond descriptive heterogeneity toward a mechanistic understanding of tumor organization, with implications for why durable control of GBM will likely require coordinated disruption of multiple progenitor programs rather than elimination of any single cell type.

## Acknowledgements

We would like to thank the members of the Bhaduri Lab for their insightful advice and comments on the study. We would like to thank the Broad Stem Cell Research Center Flow Cytometry core for their help in isolating cells for this project, and Charina Julian (UCSF) and Suhua Feng (UCLA) for help with running sequencing. We would like to thank Sergey Mareninov and others at the Brain Tumor Translational Research Core at UCLA for enabling tumor sample acquisition. We would like to thank UCLA Technology Center for Genomics and Bioinformatics (TCGB) for their help with the whole exome sequencing library preparation and sequencing. We would like to acknowledge the assistance of the Molecular Genomics Core at the Duke Molecular Physiology Institute, Duke University School of Medicine, for the generation of data for the manuscript. We would like to thank Maximilian Haeussler (UCSC) for his help in compiling the genome browser. This study was generously funded by support to AB from: Swim Across America, Jonsson Comprehensive Cancer Center, Sloan Research Fellowship from the Alfred P. Sloan Foundation, NIH NCI P50CA211015 including a Career Enhancement Program Award, The Sontag Foundation (Distinguished Scholar Award), V Scholar Award from The V Foundation, The Uncle Kory Foundation, The American Cancer Society (CSCC-Team-23-980262-01-CSCC), The Margaret Early Medical Research Trust, The Rose Hills Foundation, Broad Stem Cell Research Center, Pew Charitable Trusts, The Alexander and Margaret Stewart Trust, and the McKnight Endowment Fund for Neuroscience. Funding for EF was provided by the David Geffen Scholarship and the UCLA-Caltech Medical Scientist Training Program (T32GM152342). BW was supported by CIRM funding DISC0-14514, a collaborative grant with AB. BW was additionally supported by NIH/NIMH RF1MH132662 and NIH/NHGRI U24HG002371. This material is based upon work by JAS supported by the National Science Foundation Graduate Research Fellowship Program under Grant No. DGE-2034835. Any opinions, findings, and conclusions or recommendations expressed in this material are those of the author(s) and do not necessarily reflect the views of the National Science Foundation.

## Author Contributions

The study was conceptualized by E.F. and A.B. Experimental design was performed by E.F., A.B., D.N, and B.D. P.R.N. and B.W. assisted with bioinformatic pipeline development. E.F., D.J.A., K.Y., M.X.L., W.G., R.L.K., H.A.T., N.A.B., C.T., V.H., D.C., T.P., J.A.S., S.B., A.M., J.C., L.N.D., A.V., C.D., L.W.P., S.G.G., V.J., M.A. performed experiments and informatics. Data interpretation was performed by E.F., A.B., E.E.C, and D.R.R. Primary tissue samples were provided by K.S.P., R.G.E, M.B., and L.M.L. Analysis was performed by E.F. The manuscript was written by E.F. and A.B. with input and edits from all authors.

## Data Availability

Our GBM meta-atlas and all original datasets generated in this study are browsable and downloadable in the UCSC Genome Browser: (https://gbm-nvp.cells.ucsc.edu), in which the data are also available as Seurat objects for download. New raw data collected from this study have been deposited in dbGAP: :phs003936.v1.p1.

## Methods

### Meta-atlas dataset acquisition and uniform quality control

Gene count matrices for individual cells and corresponding metadata were obtained from various sources. For all datasets except those from Couturier and Bhaduri were downloaded from the Gene Expression Omnibus (GEO), and cells from the same individual were combined. All data not obtained from GEO were obtained through personal communication. While many of the datasets contained multiple tumor types, only direct-from-patient IDH1^WT^ primary adult GBM samples were incorporated into our analysis. The 10X-derived count matrices and directories were processed using a standard pipeline to generate Seurat objects^64^ (Seurat version 4). Normalization of the counts was performed as needed, and cells with fewer than 500 detected genes and more than 10% of UMIs mapping to mitochondrial genes were filtered out. Genes detected in fewer than three cells were omitted. Datasets were subset to contain only “tumor” cells based on the paper’s published annotations, or by cluster-based methods if published annotations were unavailable.

### Batch corrected integration for visualization and cell type annotation

For visualization purposes, we constructed the UMAP for our meta-atlas by integrating the specified datasets using conventional Seurat pipelines. Similarities between cells from different Seurat objects were identified and used as anchors to harmonize the data, employing the PCA method. FindVariableFeatures and ScaleData functions to prepare the data for principal component analysis (RunPCA function). Significant principal components were identified following the methods described by Shekhar et al.^65^. Graph-based clustering was performed using the FindNeighbors and FindClusters function. Differential gene expression analysis (FindAllMarkers) was used to identify cluster markers, reporting only genes positively enriched in a cluster. Annotation of cell identity based on integrated UMAP space was performed using a combination of annotation methods, including comparison to published datasets from GBM^10,20^, normal development^51,54,66^, and the adult brain^23,67^.

### Module scoring and normalization

Gene modules were compiled from two sources: (i) published gene lists defining transcriptional programs or cell states^24,49,50,52,67^(ii) in-house-derived gene sets generated by calculating cluster marker genes from author-annotated single-cell RNA-sequencing datasets^23,51^. For all analyses, module activity was quantified at the single-cell level using the AddModuleScore function in Seuratv5 ^68^, which computes the average expression of genes within each module relative to expression-matched control genes. To enable comparison across modules and datasets, raw module scores were z-score normalized within each dataset prior to downstream analysis.

### Cell state assignment based on module scores

For analyses requiring assignment of individual cells to a transcriptional program or cell state, cells were assigned using a winner-take-all strategy based on normalized module activity. Specifically, for a given set of mutually exclusive programs (for example, Neftel 2019 cell states or Nomura 2025 base programs), each cell was assigned to the program for which it exhibited the highest z-scored module value. This assignment was performed independently of clustering used to define meta-atlas cell types and was applied selectively for visualization, comparison, and mapping between annotation frameworks throughout the study.

### Reference-based label transfer using query-reference mapping

To compare and transfer cell type annotations across datasets, we performed reference-based label transfer using a query-reference mapping framework implemented in Seurat^68^. In each analysis, one dataset was designated as the reference and contained predefined cell type annotations, while the second dataset was treated as the query and processed independently according to dataset-specific quality control and filtering criteria prior to mapping. Shared transcriptional structure between reference and query datasets was identified by computing anchors using canonical correlation analysis-based feature alignment. Query cells were then projected into the low-dimensional embedding of the reference dataset using the MapQuery function, and reference annotations were transferred to query cells based on nearest-neighbor relationships in the shared embedding space. This approach enabled consistent annotation of query datasets without reclustering or redefining cell identities and allowed direct assessment of annotation concordance across datasets. Label transfer was performed independently for each reference-query pairing used in the study.

### Gene score correlation analysis

To quantify transcriptional similarity between cell populations across datasets, we performed gene score-based correlation analyses using cluster-level marker genes. Cluster markers were identified independently within each dataset using Seurat FindAllMarkers with parameters only.pos = TRUE and min.pct = 0.25, restricting analysis to genes positively enriched in each cluster. For each marker gene, a gene score was calculated as a composite metric reflecting both specificity and enrichment in progenitor populations. The gene score was defined as the ratio of the fraction of progenitor cells expressing the gene to the fraction of non-progenitor cells expressing the gene, multiplied by the average log_2_ fold change of expression in the progenitor population. This formulation emphasizes genes that are both selectively and strongly enriched in progenitor cells while down-weighting broadly expressed markers. Gene scores were aggregated at the cluster level to generate ranked gene score profiles for each cluster within each dataset. Pairwise correlations between clusters from different datasets were then computed using these gene score profiles to assess transcriptional similarity.

### Processing and annotation of the Nomura GBM dataset

Single-cell RNA-sequencing data from the Nomura GBM study were reprocessed with platform-aware quality control. Although the original dataset included both Smart-seq and 10x Genomics data, analyses were restricted to the 10x Genomics subset, which corresponds to the cell counts and annotations reported in the original publication. Quality filtering was applied using more stringent thresholds than those originally reported, requiring a minimum of 500 detected genes per cell. This filtering step retained approximately 600,000 high-quality cells. Cells were categorized according to the original study’s copy-number-based annotations, and only cells classified as malignant were retained; non-malignant cells and cells with ambiguous copy number status were excluded. After filtering, 241,708 malignant cells remained for downstream analyses, a modest reduction relative to the original report due to the additional quality-control criteria. Transcriptional cell types and states were assigned using reference-based mapping, in which malignant cells were projected onto a curated GBM meta-atlas using Seurat MapQuery^68^ to ensure consistent cell state annotation across datasets.

### Immunofluorescent staining

Human primary GBM samples were obtained from patients undergoing surgery at Ronald Regan UCLA Medical Center. All patients signed informed consent forms for tissue collection under institutional IRB#10-000655. Samples were fixed in 4% paraformaldehyde for overnight at 4°C, rinsed with PBS, and equilibrated in 30% sucrose in PBS for 24-48 hours at 4°C. Post-equilibration, tumors were embedded in a 1:1 mixture of OCT and 30% sucrose, then frozen on dry ice. Frozen blocks were either stored at -80°C or processed on the cryostat into 10-16 μm-thick sections for immunofluorescence staining. Sections were first rinsed with PBS for 15 minutes and subjected to antigen unmasking using a citrate-based solution (10 mM sodium citrate, pH 6) heated to 95°C for 20 minutes. Following this, sections were permeabilized and blocked with a buffer containing 5% donkey serum, 3% bovine serum albumin, and 0.1% Triton X-100 in PBS for 30 minutes at room temperature. Primary antibodies were incubated overnight at 4°C in the blocking buffer: rabbit anti-PDGFRβ (1:100), goat anti-PDGFRβ (1:100), mouse anti-Nestin (1:500), mouse anti-NOTCH3 (1:200), rabbit anti-COL1A1 (1:200), and rabbit anti-HOPX (1:500). Following primary antibody incubation, sections were washed three times with PBS for 10 minutes each. Secondary antibody incubations, including DAPI (1:1000), were performed in blocking buffer for 2 hours at room temperature using the following secondary antibodies: AlexaFluor 555 donkey anti-goat (1:500), AlexaFluor 555 donkey anti-mouse (1:500), and AlexaFluor 488 donkey anti-rabbit (1:500). Slides were then mounted with ProLong Gold antifade reagent and stored at 4°C for imaging using the ZEISS LSM 880 confocal system or EVOS M5000 digital-inverted benchtop microscope.

### Analysis of spatial transcriptomics data: Visium

Visium Seurat objects from Tsyben et al.^46^ were downloaded and spatial spot identities were assigned using RCTD (spacexr) with the GBM meta-atlas as the single-cell reference. The RCTD reference was generated from meta-atlas raw RNA counts and corresponding cell type annotations. Visium sections were merged by concatenating spot centroid coordinates and aligning per-slice count matrices prior to construction of the SpatialRNA object. RCTD was run in doublet mode (doublet_mode = "doublet"), and spot identities were assigned using the primary classification (first_type), which was appended to the Visium Seurat object for downstream analysis and visualization.

### Xenium ISS data collection

For Xenium data collection, deparaffinization and rehydration of tissue slices were performed by immersing the FFPE sections in xylene and an ethanol gradient of decreasing concentrations to prepare the sections for the Xenium v1 procedure. Spatial transcriptome data were collected using the Xenium Analyzer platform. Oligonucleotide probes complementary to target mRNAs were synthesized and labelled with unique fluorescent tags. Tissue sections were incubated with these probes under optimal conditions to ensure specific binding. High-resolution fluorescence microscopy was used to acquire images of the tissue sections. Multiple imaging rounds were performed to detect various fluorescence signals corresponding to different transcripts. The images were then stitched together to reconstruct the complete spatial map of the tissue section.

### Re-segmentation of Xenium ISS data

To restrict transcript assignment to nuclear regions of cells, we reprocessed the initial Xenium data outputs using Xenium Ranger v1.6 with modified segmentation parameters that limited cell boundaries to the detected nuclei only. In this re-segmentation step, the nuclear expansion distance was explicitly set to 0 um, preventing outward expansion beyond the DAPI-based nuclear mask. The resulting nuclei-restricted segmentation outputs were used for all downstream analyses. This nuclei-focused approach provided a more conservative and precise transcript assignment strategy, critical for analyses driven by cell identity and transcriptional state in complex brain tumor tissue.

### Multiome sequencing and analysis

Multiome sequencing was performed on nuclei isolated from flash frozen tumors. Nuclei isolation was performed using the 10X demonstrated protocol for Single Cell Multiome ATAC + Gene Expression Sequencing. Briefly, flash frozen tissue is used frozen and immersed in 500uL of 0.1X Lysis Buffer. It is incubated on ice for 5 minutes, pipette mixed until dissolution of tissue using a 1000 uL wide bore tip, and incubated on ice for another 10 minutes. 1 mL of Wash Buffer is added to the lysed cells and mixed, before centrifuging at 4°C at 500 rcf. The supernatant is removed and is washed twice more, before nuclei are counted and spun again at 4°C at 500 rcf before resuspension in nuclei isolation buffer. Buffer components are as below:

Lysis Buffer (1X):

Tris-HCl (pH 7.4) - 10 mM
NaCl - 10 mM
MgCl2 - 3 mM
Tween 20 - 0.1%
Nonidet P40 Substitute - 0.1%
Digitonin - 0.01%
BSA - 1%
DTT - 1mM
RNase inhibitor 40 U/µl - 1 U/µl
Brought to volume in nuclease-free water

Lysis Dilution Buffer (1X):

Tris-HCl (pH 7.4) - 10 mM
NaCl - 10 mM
MgCl2 - 3 mM
BSA - 1% DTT - 1mM
RNase inhibitor 40 U/µl - 1 U/µl
Brought to volume in nuclease-free water

Wash Buffer

Tris-HCl (pH 7.4) - 10 mM
NaCl - 10 mM
MgCl2 - 3 mM
Tween 20 - 0.1%
BSA - 1%
DTT - 1mM
RNase inhibitor 40 U/µl - 1 U/µl
Brought to volume in nuclease-free water

Nuclei Resuspension Buffer

10X Genomics provided nuclei resuspension buffer (20X)
DDT - 1mM
RNase inhibitor 40 U/µl - 1 U/µl
Brought to volume in nuclease-free water

Multiome capture was performed by 10X Genomics manufacturer’s protocol. Libraries were generated by this protocol and sequencing on the NextSeq 2000. Alignment was performed with cellranger-arc. Quality control for RNA and ATAC was performed using Signac default parameters^69^. For the cell type designation, MapQuery from Seurat v5 was used to map meta-atlas cell types onto the multiome transcriptional data. CNV at the single-cell level was performed using epiAneufinder^47^. These annotations were applied to the results from epiAneufinder and plotted using Morpheus to generate heatmaps for the full analysis and for the NVP populations alone. Chromosome level deviations from diploid were calculated using metrics from the IGV Browser and the Mann-Whitney statistical test, as is common in the field^70^.

### Mutation and clone analysis based on multiome data

Somatic mutations were inferred from the ATAC output of single-cell multiome data aligned with CellRanger ATAC (v2.0.0) to the GRCh38 reference genome. BAM files were processed with Picard Tools (SortSam, MarkDuplicates) and indexed with Samtools. Per-barcode BAM files were generated and restricted to cell barcodes that passed quality control (QC) based on RNA (>500 counts) and ATAC (>300 fragments) metrics in a Seurat object constructed with Signac. Mutect2 and FilterMutectCalls (GATK v4.2.2.0) were applied to each barcode BAM using the GRCh38 reference genome to identify somatic variants. Variants were retained if labeled as “PASS” and located on chromosomes 1-22. To enrich for coding mutations, the merged VCF was intersected with coding exon annotations obtained from the UCSC Genome Browser. A binary mutation matrix was constructed and clone calling was performed using a Jaccard correlation–based strategy. Barcodes annotated as Mixed_Vascular (NVP) via MapQuery projection to the GBM meta-atlas were used to identify mutations shared between NVP and non-NVP cells. A whitelist of these shared mutations was applied to the full binary matrix. Pairwise Jaccard correlations were computed, and a correlation threshold was selected based on the empirical distribution. Groups of cells with mutual similarity above this threshold were assigned to the same clone.

### Tumor dissociation

Primary tumors were obtained from Ronald Reagan Hospital at UCLA, with patient consent and Institutional Review Board approval (IRB #10-000655 and #21-000108). Tumors were dissected into small fragments using a sterile scalpel and transferred to 5 mL microcentrifuge tubes containing 2.5 mL Papain and 125 μL DNase. Samples were incubated at 37°C for 45-60 minutes, with vigorous shaking by hand for 10 seconds every 5 minutes to facilitate dissociation. Post-incubation, the tissue was further dissociated by trituration and centrifuged at 300 x g for 5 minutes. The cell suspension was then passed through a 40 μm filter to remove debris and counted. For further debris removal, cells were subjected to an ovomucoid density gradient according to the Papain Tissue Dissociation Kit instructions. Briefly, up to 20 million tumor cells were resuspended in a solution of ovomucoid inhibitor, DNase, and EBSS and carefully layered on top of an ovomucoid inhibitor cushion. The gradients were centrifuged at 70 x g for 6 minutes. The supernatant was discarded, and the tumor cells were collected, combined, and resuspended in media or FACS buffer for subsequent analyses.

### Fluorescence-activated cell sorting for PDGFRB+ enrichment

The cell suspension was incubated on ice for 30 minutes with Phycoerythrin (PE)-conjugated mouse anti-hPDGFRβ (1:100) and DAPI (1:2000) in FACS buffer (1% BSA, .1% Glucose in HBSS). Following incubation, cells were washed and sorted using a Sony SH800, BD FACSAria, or Bio-Rad S3e cell sorter. All fluorescence-activated cell sorting (FACS) gates were set using unlabeled cells and single-color controls to ensure accurate gating and minimize background fluorescence. PDGFRβ+, DAPI-cells were collected into Sasai3 media.

### Single cell capture and sequencing

Single cells, either sorted by FACS or derived from dissociated tumor cells, were captured using the 10X Genomics Chromium v3.1 3’ capture protocol on the 10X Chromium system. Targeting 10,000 cells for capture, all cells were used if fewer than 20,000 cells were retrieved post-sorting. The capture process was conducted according to the manufacturer’s instructions, and library preparation adhered to their guidelines. Sequencing was performed on an Illumina NovaSeq 6000 or NovaSeq X.

### Single-cell analysis and quality control

Single-cell RNA sequencing (scRNA-seq) reads were aligned to a custom reference using a species-specific reference genome. For human samples, data was aligned to the GRCh38 human reference genome, incorporating sequences for the CellTag UTR and GFP.CDS. For mouse samples, data was aligned to the mm10 mouse reference genome, incorporating sequences for eGFP using CellRanger mkref. Cell-by-gene count matrices were generated using the 10X Genomics CellRanger pipeline with default parameters. Subsequent data analysis was performed using the Seurat R package in R Studio. Single-nucleus samples were run through CellBender^71^, a package capable of removing background noise associated with ambient RNA in snRNA-seq data. Cells expressing a minimum of 500 genes and exhibiting less than 10% mitochondrial gene content were retained. Genes expressed in at least three cells were also retained. Unique Molecular Identifier (UMI) counts were normalized via logarithmic transformation with a scaling factor of 10,000. Principal Component Analysis (PCA) was conducted on the scaled data using the top 2,000 variable genes. The number of dimensions (dims) was determined according to previously described methods^65^). Specifically, significant principal components (PCs) were selected based on the larger value between the square of the standard deviation of PCA scores (Seurat.Obj@reductions$pca@stdev^2) and the square root of the ratio of the number of genes to the number of cells plus one (sqrt(length(row.names(Seurat.Obj)) / length(colnames(Seurat.Obj))) + 1)^2. Cells were clustered in the PCA space using Seurat’s FindNeighbors and FindClusters functions with a resolution parameter set to 2.0. Visualization of cell clusters was performed using Uniform Manifold Approximation and Projection (UMAP) based on the previously defined dimensions. Doublets were predicted and excluded using the DoubletFinder R package^72^ with default parameters.

### Copy number variation analysis

To reduce the risk of false positives or negatives in tumor cell identification via FACS, copy number variation (CNV) analysis was conducted using InferCNV^73^ with default settings. Batch-matched naïve organoids were used as reference datasets for HOTT experiments. Other reference datasets were used as indicated in the figure legends. Where appropriate tumor, datasets were subset to contain tumor cells only.

### Matrigel tube formation assay

Tumor cells were dissociated using the Papain Dissociation System (Worthington) and stained with fluorophore-conjugated antibodies: CD31-FITC (1:200), PDGFRβ-PE (1:200), CD45-BUV395 (1:200), and DAPI (1:400). Fluorescence-activated cell sorting (FACS) was performed on a BD FACSAria. Cells were first gated by size (FSC-A vs. SSC-A), then singlets were selected by forward scatter (FSC-H vs. FSC-A), followed by gating of live cells (DAPI-). Populations were defined as follows: immune cells (CD31-CD45⁺), endothelial cells (CD45-CD31⁺), and PDGFRβ-enriched (CD45-CD31-PDGFRβ⁺). Cells were collected in EGM-2 medium (Lonza) and plated on undiluted Growth Factor-Reduced Matrigel (Corning) in 96-well plates pre-coated with Matrigel. Cultures were maintained in EGM-2 for 7 days with media changes every other day. Tube formation was assessed daily using brightfield microscopy (Echo REVOLVE).

### Stem cell culture for cortical organoid generation

The human embryonic stem cells (hESCs) UCLA6 was cultured as previously described^74–76^. Stem cells were maintained on Matrigel-coated 6-well plates in mTeSR Plus medium supplemented with 10% mTeSR Plus Supplement and 1X Penicillin/Streptomycin or Primocin. Media was changed every other day, and cells were passaged upon reaching >75% confluence. For passaging, ReLeSR was applied at room temperature for 1 minute, aspirated, and cells were incubated at 37°C for 5 minutes. Cells were then dissociated into smaller clusters and replated at a 1:4 or 1:6 ratio on new Matrigel-coated plates. For cryopreservation, the passaging procedure was followed until the final step, where cells were resuspended in 1 ml of mFreSR per well. The cell suspension was transferred to cryovials and stored at -80°C for 24-48 hours before transfer to liquid nitrogen for long-term storage.

### Cortical organoid generation

Cortical organoids were generated following an adapted protocol from Kadoshima et al.^77^, in alignment with other studies^74–76^. Briefly, human embryonic stem cells (hESCs) at >75% confluence in 6-well plates were treated with 1 mL Accutase per well and incubated at 37°C for 5 minutes. Cells were then washed with 1 mL of Sasai1, composed of GMEM, 20% KnockOut Serum, 0.1 mM β-mercaptoethanol, 1X Non-Essential Amino Acids (NEAA), 1X Sodium Pyruvate, and 1X Penicillin/Streptomycin or Primocin. Cells were detached by scraping and collected in 15 mL tubes, followed by centrifugation at 300 x g for 5 minutes. The cell pellets were resuspended in Sasai1 supplemented with 20 μM Y27632 (Rock inhibitor), 5 μM SB431542 (TGF-β inhibitor), and 3 μM IWR1-endo (Wnt signaling inhibitor). A total of 1 million cells in 10 mL of Sasai1 with the small molecules were seeded into 96-well V-shaped low attachment plates to aggregate over 72 hours. On Day 3, 50 μL of medium was replaced with 100 μL of fresh Sasai1 containing the small molecules. Media changes were performed every other day until Day 7, when the Rock inhibitor was omitted. On Day 18, organoids were transferred to ultra low attachment 6-well plates with media changes every other day. From Day 18 to Day 35, Sasai 2, consisting of DMEM/F-12 with Glutamax, 1X N-2 supplement, 1X Lipid Concentrate, and 1X Penicillin/Streptomycin or Primocin, was used. From Day 35 onward, Sasai3 was applied, containing DMEM/F-12 with Glutamax, 1X N-2 supplement, 1X Lipid Concentrate, 1X Penicillin/Streptomycin or Primocin, 10% Fetal Bovine Serum, 5 μg/mL Heparin, and 0.5% Growth factor-reduced Matrigel. Throughout the culture period, live images were periodically taken to monitor organoid growth, and immunostaining was performed at Weeks 5 and 8 to confirm neuronal differentiation.

### Transduction with CellTag and transplantation onto cortical organoids

Lentivirus containing the CellTag-V1 barcode library was either directly purchased from Addgene or manufactured by a commercial vendor (Vectorbuilder) using the DNA plasmid library purchased from Addgene. For commercial generation of CellTag virus, the libraries were sequenced using MiSeq and CellTags were filtered in order to create a whitelist based on the standard CellTag workflow. The Vectorbuilder lentiviral library displayed a complexity of 7,989 barcodes, while the Addgene lentiviral library had a complexity of 19.974 barcodes. Appropriate whitelists were applied to samples during data processing depending on which lentivirus was used in the experiment. Immediately following FACS, cells were resuspended in Sasai3 medium. CellTag virus and polybrene (1:1000) were added to the cells, which were then incubated at 37°C for 60 minutes with gentle rotation. Amount of CellTag virus was based on a target MOI of 4 using this formula: TUtotal = (MOI x Cell Number)/Viral titer (TU/μl). Post-transduction, cells were washed three times with warm PBS and Sasai3 medium. The transduced tumor cells were resuspended in 20 μL of Sasai3 medium and transplanted onto cortical organoids using the hanging drop method. For the hanging drop method, 8-12-week-old human cortical organoids were transferred to the lid of a 10cm dish using wide-bore 1000 μL tips. Excess media was removed, and 10-15 μL of the tumor single-cell suspension was added atop each organoid. The lid was then carefully inverted onto a 10 cm dish containing 10 mL of base culture medium to prevent evaporation during the hanging drop stage. These hanging drop co-cultures were maintained at 37°C for 12-16 hours, after which they were transferred to ultra-low attachment 6-well plates with Sasai3. Tumor cells typically surrounded the organoids and began migrating inward after a few days. These co-cultures were maintained for 12 - 18 days, with media changes performed three times per week, before being harvested for analysis.

### Organoid-tumor transplant dissociation and collection of GFP+ cells by FACS

Tumor-transplanted organoids were transferred to a 1.7 mL Eppendorf tube containing 1 mL Papain and 50 μL DNase, and incubated at 37°C for 45-60 minutes. During the initial 10 minutes of incubation, the tube was shaken vigorously by hand for 10 seconds to aid dissociation, a process repeated every 5 minutes. Following incubation, the material was further dissociated by trituration and centrifuged at 300 x g for 5 minutes. The resulting pellet was resuspended in cold FACS buffer, filtered through a 40 μm mesh, and stained with DAPI to identify non-viable cells. DAPI-/GFP+ tumor cells were sorted and collected for downstream single-cell capture. Flow cytometry and cell sorting were performed using a Bio-Rad S3e. Gates were set to remove debris, eliminate doublets, and exclude dead cells using DAPI staining. Gates for GFP positivity were calibrated using control, batch-matched organoids without transplantation.

### Side library preparation for CellTag-barcode association

To enable direct pairing of 10x Chromium cell barcodes with corresponding CellTag sequences, a targeted auxiliary library was constructed using genomic DNA derived from the same 10x Chromium reactions processed for gene expression analysis. This side library was selectively amplified using primers engineered to both capture the CellTag cassette located downstream of the GFP reporter and incorporate Illumina-compatible sequencing adapters.

Primer sequences (5′→3′) were as follows:

Forward (prtlTruSeqR1_1): ACACTCTTTCCCTACACGACG
Reverse (TruSeqR2-eGFP-3′): GTGACTGGAGTTCAGACGTGTGCTCTTCCGATCTggcatggacgagctgtacaag

The reverse primer was designed to anneal downstream of the GFP coding region, thereby ensuring amplification of the CellTag cassette physically linked to the 10x cell barcode. Library amplification was performed using a high-fidelity DNA polymerase to append TruSeq-compatible adapter sequences. A subsequent indexing PCR was carried out to introduce sample-specific indices and complete Illumina adapter structures. Final libraries were subjected to size selection and purification using SPRI bead-based cleanup to enrich fragments corresponding to the expected amplicon length.

### Generation of a custom reference for side library alignment

Reads from the side library were mapped to a custom CellTag reference generated using Cell Ranger (v8.0.0) via the cellranger mkref workflow. This reference was composed of consensus CellTag sequences derived from the whitelisted CellTag library, each concatenated to a constant backbone sequence containing the flanking reporter and adapter regions. The full reference sequence was structured as follows:

GGCATGGACGAGCTGTACAAGTAAACCGGT[CellTag] GAATTCGATGACAGGCGCAGCTTCCGAGGGATTTGAGATCCAGACATGATAAGATACATTGATGAGTTTGGACAAACCAAAACTAGAATGCAGTGAAAAAAATGCCTTATTTG

Prior to reference construction, individual CellTag sequences were processed to generate consensus sequences, thereby collapsing sequencing errors and reducing barcode redundancy. The resulting FASTA and corresponding annotation files were assembled into a Cell Ranger-compatible reference using cellranger mkref. Side library reads were subsequently aligned and quantified using standard Cell Ranger alignment pipelines, configured to support targeted recovery of CellTag-barcode associations.

### CellTag processing and clone inference from side libraries

CellTag-based lineage inference was carried out using sequencing data generated from a dedicated side library, which was processed separately from the gene expression libraries. Clone calling was limited to the population of tumor singlets that passed quality control in the gene expression analysis, ensuring that inferred lineage relationships were restricted to high-confidence malignant cells. Within this QC-filtered subset, cells were required to contain between 1 and 20 detectable CellTags to be considered eligible for clone inference. Cells falling below this range were excluded due to insufficient barcode complexity, whereas cells exceeding this threshold were removed to avoid inclusion of potential technical artifacts associated with abnormally high barcode counts. For eligible cells, CellTag counts were binarized, and the resulting binary matrices were used to compute pairwise Jaccard similarity scores on a per-sample basis. Clonal relationships were defined by applying a Jaccard similarity threshold of 0.7, with connected components in the resulting similarity graph designated as individual clones. The overall CellTag similarity calculation and clone-calling strategy were adapted from established barcode-based lineage reconstruction approaches^56^. Cells that did not meet the criteria for clone inference were retained in the dataset but annotated as non-contributing to clone calling. Clone identifiers were subsequently merged into the gene expression metadata for all QC-passing cells, enabling downstream analyses to incorporate clonal information without altering the underlying gene expression quantification.

### Clone-level cell type co-occurrence and enrichment analysis

Clone-level relationships between transcriptionally defined GBM cell types were quantified using CellTag-derived clonal identities. For each clone, a binary clone × cell type presence matrix was constructed, in which a given cell type was scored as present if at least one cell of that type was detected within the clone, independent of cell number. Clones lacking valid CellTag assignments or cell type annotations were excluded from analysis, and Low.Quality annotations were removed. Observed co-occurrence between pairs of cell types was defined as the number of clones in which both cell types were present. Expected co-occurrence was calculated under a null model assuming independence, using background cell type frequencies computed across all annotated cells in the dataset and scaled by the total number of clones. Enrichment of co-occurrence was quantified as the log₂ ratio of observed to expected values, with a small pseudocount added to both terms for numerical stability.

### PDGFRβ⁺ enrichment and depletion

Dissociated GBM cells were stained for CD45-APC (1:20) and PDGFRβ-PE (1:20) and DAPI (1:2000) before FACS analysis. Cells were first gated by size (FSC-H vs. SSC-H), followed by singlet gating (FSC-A vs. FSC-H and SSC-A vs. SSC-H). Live cells were identified using a DAPI⁻ gate. To exclude immune cells, CD45⁻ gating was applied. The CD45⁻PDGFRβ⁺ fraction was isolated as the enriched (POS) population, while CD45⁻PDGFRβ⁻ cells were collected as the depleted (NEG) population. POS and NEG fractions were independently infected with CellTag lentivirus (as described above), introduced into the Human Organoid Tumor Transplant (HOTT) system, and subjected to single-cell RNA-seq analysis.

### Cell-Cell Communication Analysis with CellChat

The CellChat R package^78^ was employed to infer and visualize cell-cell communication networks following its standard protocol (https://github.com/sqjin/CellChat). Analyses were independently performed on PDGFRβ⁺ (POS) and PDGFRβ⁻ (NEG) tumor-derived samples using their respective Seurat objects as input. Communication probability was calculated across annotated cell types, and inferred signaling networks were stratified by interaction category, including secreted signaling, ECM-receptor, and cell-cell contact pathways. Visualization of aggregated signaling networks was performed with netVisual_aggregate, while pathway-specific interaction contributions were assessed using netAnalysis_contribution. Category-specific signaling role heatmaps were generated with netAnalysis_signaling and Role_heatmap. For pathways of therapeutic interest, directional bubble plots were constructed to examine ligand-receptor interactions originating from NVP cells toward other tumor cell types.

### Reactome Pathway and Drug Target Enrichment Analysis

Pathway and functional enrichment analysis was performed on the NVP gene signature using Enrichr, including the Reactome and MSigDB databases. The top 20 significantly enriched pathways were selected based on adjusted p-value rankings. Genes contributing to each pathway were extracted, and those that appeared across multiple pathways were retained for further evaluation. Genes known to be associated with toxicity or essential physiological roles (e.g., platelet adhesion, extracellular matrix function) were excluded from downstream analysis. To assess therapeutic relevance, the filtered gene list was queried against the Drug Gene Interaction Database (DGIdB) to identify existing drug-target associations, with prioritization given to FDA-approved compounds. Genes lacking approved inhibitors but enriched across multiple significant pathways were retained as candidates for future investigation.

### sgRNA candidate selection

Candidates for *in vivo* knockout were identified by intersecting human and mouse NVP signatures. The mouse signature was derived calculating cluster markers from clusters in the mouse single-cell dataset^58^, annotated as “NVP” following projection onto the GBM meta-atlas. These mouse cluster markers were intersected with those calculated from the human NVP-enriched cluster (Figure 3). Feature plots for all intersecting genes were generated, prioritizing genes specific to the putative NVP population in both mouse and human datasets. Additionally, genes with transcription factor functions were preferred due to their potential role in regulating critical gene programs within our clusters of interest. Sox18 and Foxc1 were selected based on these criteria. In addition to transcription factors, intersecting and specific genes that represent markers of mixed vascular and neural progenitor identities were also considered as candidates. Fam107a (outer radial glia), Cldn5 (endothelial), and Rgs5 (mural) were chosen for their representative roles in these cellular identities.

### sgRNA design

For the design of single-guide RNAs (sgRNAs), we employed the CRISPR design tool available at CRISPOR^79^ (http://crispor.tefor.net/). sgRNA sequences were designed against the mm10 genome, incorporating a 20bp-NGG sequence as the SpCas9 PAM site. Genomic sequences for the genes of interest were sourced from the NCBI database. Target sites were selected based on the following criteria: (i) the presence of a protospacer adjacent motif (PAM) sequence (NGG) required for SpCas9 binding, (ii) positioning within the coding region of the gene, preferably targeting early exons to enhance the likelihood of generating loss-of-function alleles, and (iii) minimal off-target potential as predicted by the CRISPR design tool, prioritizing sgRNAs with high on-target activity scores and low off-target scores. For genes with only a single exon (Foxc1 and Cldn5), we designed two sgRNAs: one at the 5’ end and one at the 3’ end to maximize the chances of completely disrupting protein expression. For other targets, sgRNAs were designed against the second exon to achieve efficient gene disruption.

### Cloning of an all-in-one CRISPR/Cas9 vector expressing multiple gRNAs

The cloning of an all-in-one CRISPR/Cas9 vector expressing multiple gRNAs was conducted using the Multiplex CRISPR/Cas9 Assembly system available from Addgene^80^. This approach utilizes a Golden Gate assembly system to sequentially assemble multiple inserts into a single plasmid. The final experimental plasmid contained sgRNAs targeting Sox18, the 5’ end of Foxc1, the 3’ end of Foxc1, the 5’ end of Cldn5, the 3’ end of Cldn5, Fam107a, and Rgs5.

### In utero electroporation

Mouse gliomas were generated using the CD1 IGS mouse background, following previously described protocols^58,59^. *In utero* electroporation was performed on embryonic day 15 with a plasmid cocktail containing constructs for CRISPR-Cas9-mediated knockout of tumor suppressor genes *Nf1*, *Trp53* (*p53*), and *Pten*, referred to as the triple CRISPR (3xCr) construct. The control group received a plasmid cocktail consisting of 1.5 µg/µl *3xCr*, 2 µg/µl pGlast-PBase, and 0.25 µg/µl PBCAG-EGFP. The experimental group, termed *NVP^null^*, received an additional plasmid (also at 1.5 µg/µl) containing seven sgRNAs against NVP target genes. Mice were housed in a controlled environment with food and water available ad libitum, under a 12-hour light/dark cycle, at 20-22°C and 40-60% humidity. Both female and male mice were included in all experiments. All procedures were approved by the Institutional Animal Care and Use Committee (IACUC) at Baylor College of Medicine and conform to the US Public Health Service Policy on Human Care and Use of Laboratory Animals.

### Single-nucleus capture and sequencing of mouse tumors

At postnatal day 70 (P70), mice were euthanized, and tumors were dissected under a GFP fluorescent microscope. GFP-positive tumors were snap-frozen in liquid nitrogen for storage. Nuclei were isolated using the 10X Genomics Chromium Nuclei Isolation Kit with an 8-minute lysis time. Samples underwent two additional washing steps before proceeding to GEM capture, targeting the recovery of 10,000 cells. After standard QC, including Cellbender to remove technical artifacts associated with ambient RNA, data was processed using our Seurat pipeline and cells were annotated by projection onto the human GBM meta-atlas. “Neuronal” and “Immune” cells were excluded from downstream composition analysis, as these populations had low percentages of GFP positivity.

### Co-localization analysis using ilastik and SynBot

Images were thresholded and segmented using a machine-learning based workflow in ilastik43 (version 1.4.1b23; Pixel Classification mode). For this, a classifier was trained by manually annotating true and background signals in a training set of five randomly selected images for each channel (GFP and RFP). All available image features, including color/intensity, edge, and texture, were used as parameters for classification. The classifier was then applied to the entire dataset to generate binary masks for each channel. Segmented images were then analyzed for 2-channel (GFP and RFP) colocalization using SynBot^81^ (https://github.com/Eroglu-Lab/Syn_Bot/blob/main/Syn_Bot.ijm). Whole images were used as regions of interest, and “Circular Approximation” was selected as the analysis type to quantify the number of colocalized GFP- and RFP-positive signals in each image.

### Whole exome sequencing for sgRNA specificity

Genomic DNA was extracted from flash-frozen mouse tumors collected at P70 and at terminal timepoints using the DNeasy Blood & Tissue Kit (Qiagen). Library preparation was performed using the KAPA HyperExome V3 kit, and samples were sequenced with 2×150 bp paired-end reads on an Illumina NovaSeq X Plus 10B. Reads were aligned to the mm10 reference genome (UCSC) using BWA-MEM. Resulting BAM files were sorted, indexed, and processed with Samtools and Picard Tools. Somatic variant calling was performed using GATK Mutect2, with control tumor samples designated as matched normal references for each experimental tumor. FilterMutectCalls was applied to retain only high-confidence PASS variants on chromosomes 1-22. To assess CRISPR-Cas9 specificity, predicted Cas9 cut sites were obtained from CRISPOR. On-target loci were defined as the intended sgRNA targets, while the top five highest-scoring predicted off-target loci per sgRNA were also included. BED files representing these sites were intersected with filtered VCFs using Bedtools to quantify the number of mutations at each locus. Variant counts were summarized per target class (on-target vs. off-target) and per gene to evaluate editing fidelity across experimental tumors.

---

**Supplementary Figure 1:**
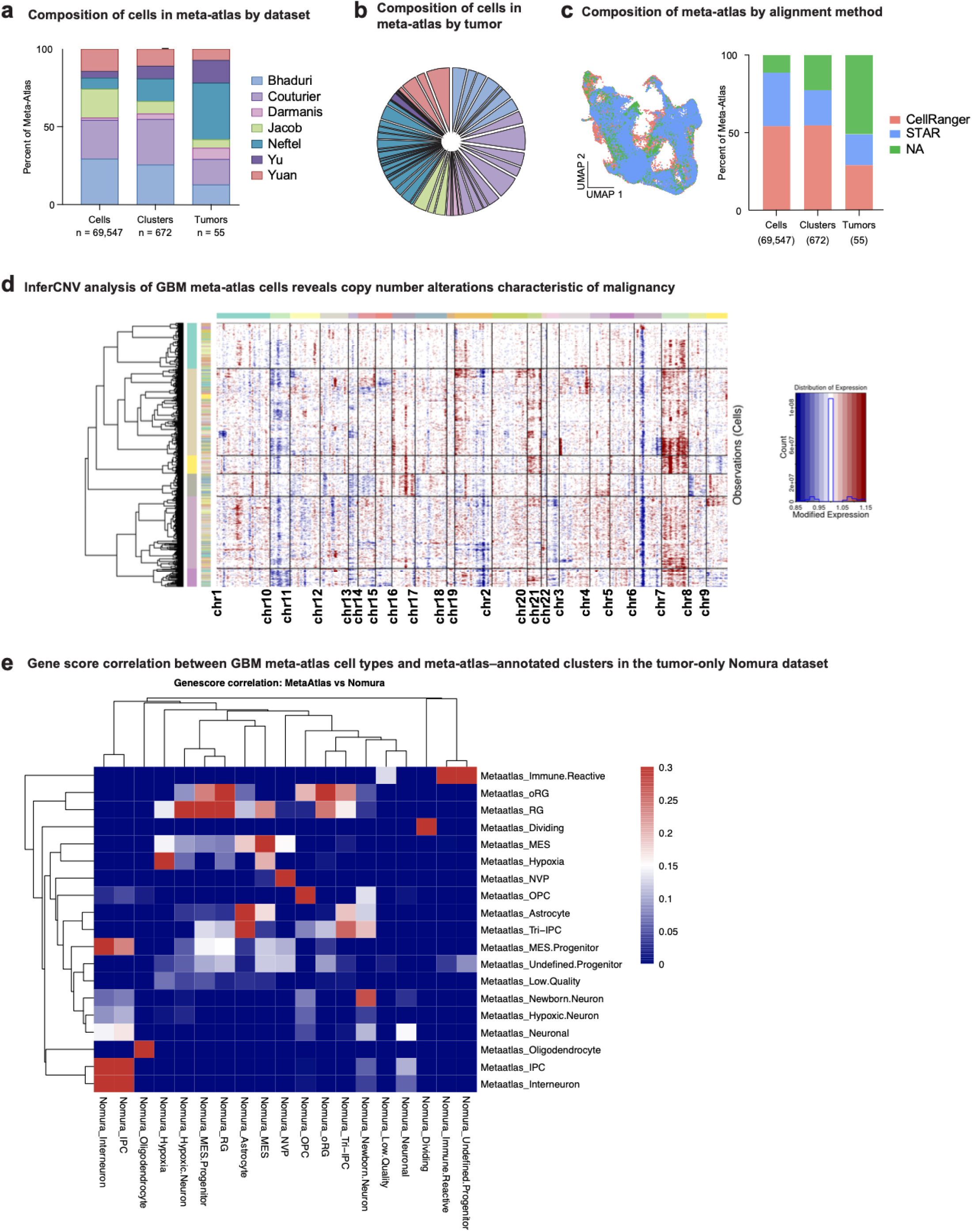
Overview and validation of the GBM meta-atlas. a) Composition of cells (n = 69,547), clusters (n = 672), and tumors (n = 55) in the meta-atlas, categorized by dataset. b) Pie chart showing the distribution of cells in the GBM meta-atlas by tumor of origin, with segments colored by contributing dataset. c) UMAP visualization of the GBM meta-atlas colored by read alignment method, including CellRanger, STAR, or not available (NA). NA denotes samples for which alignment information was not explicitly reported in the original publication. d) Heatmap showing inferred copy number variations (CNVs) across cells in the GBM meta-atlas. Copy number gains and losses are indicated in red and blue, respectively, relative to a reference composed of non-tumor cells. To construct the normal reference, a Seurat object was generated from cells annotated as non-tumor in the original meta-atlas source publications. Cells are hierarchically clustered based on inferred CNV profiles, and genomic position is shown along the x-axis, ordered by chromosome. The presence of canonical GBM-associated CNV patterns, including chromosome-wide gains and losses, supports the malignant identity of cells included in the meta-atlas. (Methods) e) Heatmap^82^ showing Pearson correlations between cluster-level gene score profiles (Methods) derived from the GBM meta-atlas (rows) and from the tumor-only Nomura dataset after projection of meta-atlas annotations (columns). High correlation along corresponding identities indicates concordant transcriptional programs across datasets and supports robust transfer of meta-atlas cell type definitions, including a specific correspondence for the NVP.

**Supplementary Figure 2.**
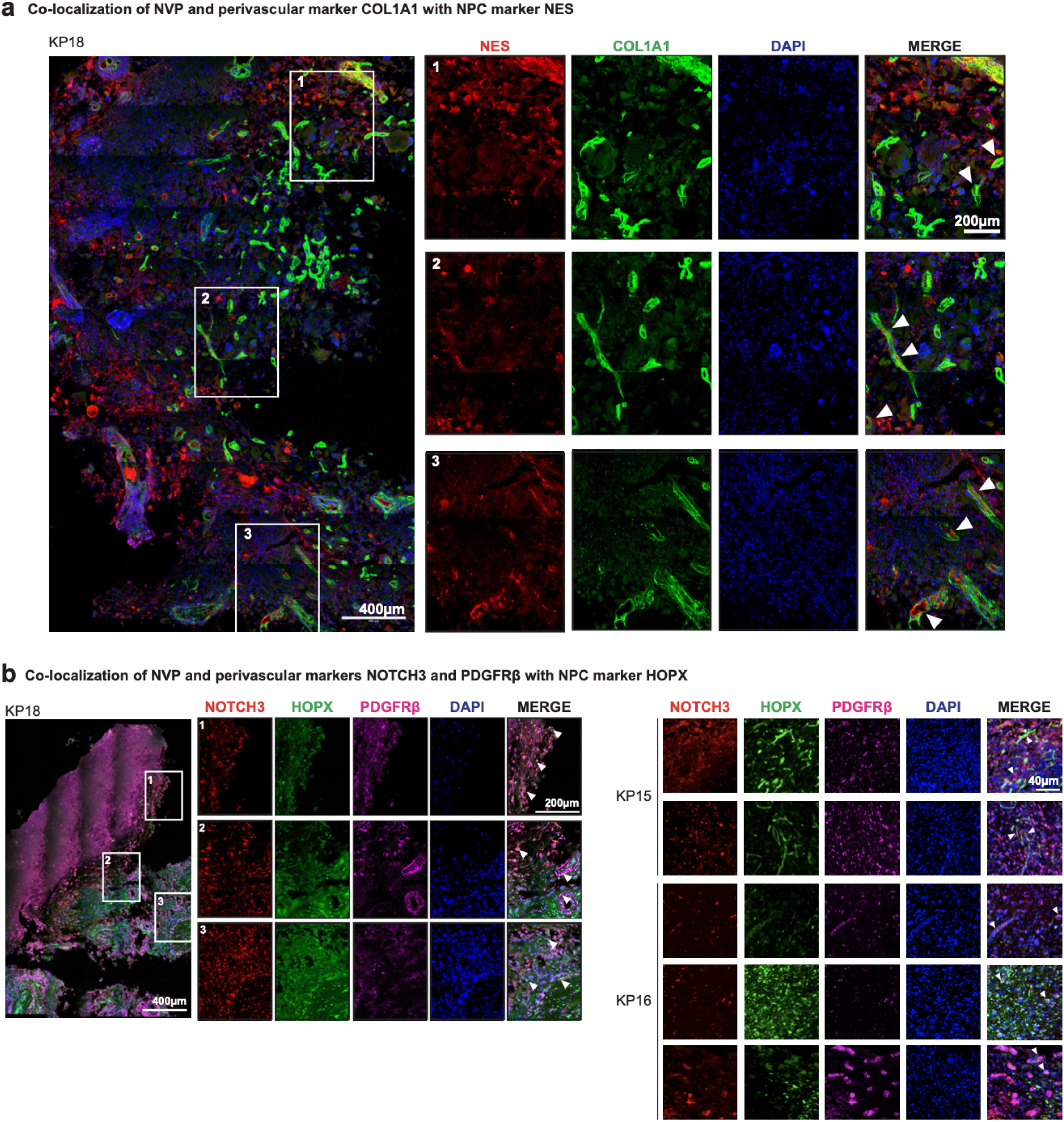
*In situ* validation of additional perivascular and neural progenitor marker co-expression in GBM. a) *In situ* immunofluorescence was performed to validate co-expression of additional perivascular and neural progenitor markers in primary GBM tissue. Left panel: 20× tile scan from tumor KP18 showing co-localization of additional perivascular marker and NVP marker COL1A1 (green) with the neural progenitor cell (NPC) marker NES (red); nuclei are counterstained with DAPI (blue). Right panels: Insets 1-3 correspond to boxed regions in the tile scan and highlight regions of marker co-expression (white arrows) at higher magnification. Co-expression is primarily localized to vascular morphologies but is also observed in tumor parenchyma without overt vascular association. Scale bar in overview image = 400 μm; scale bars in inset panels = 200 μm (1.7× zoom). Staining was performed in three tumors. Representative images are shown. b) Immunostaining in additional tumor samples demonstrates co-localization of NVP and perivascular markers NOTCH3 (red) and PDGFRβ (magenta) with the neural progenitor marker HOPX (green); nuclei are counterstained with DAPI (blue). Left panel: 20× tile scan from tumor KP18 with three insets (1-3) highlighting regions of co-expression, denoted by arrows in the merged images. Scale bar in overview image = 400 μm; scale bars in inset panels = 200 μm (1.7× zoom) Right panel: 20X images from tumors KP15 and KP16 demonstrate consistent co-localization of NOTCH3, PDGFRβ, and HOPX. Scale bar = 40 μm.

**Supplementary Figure 3.**
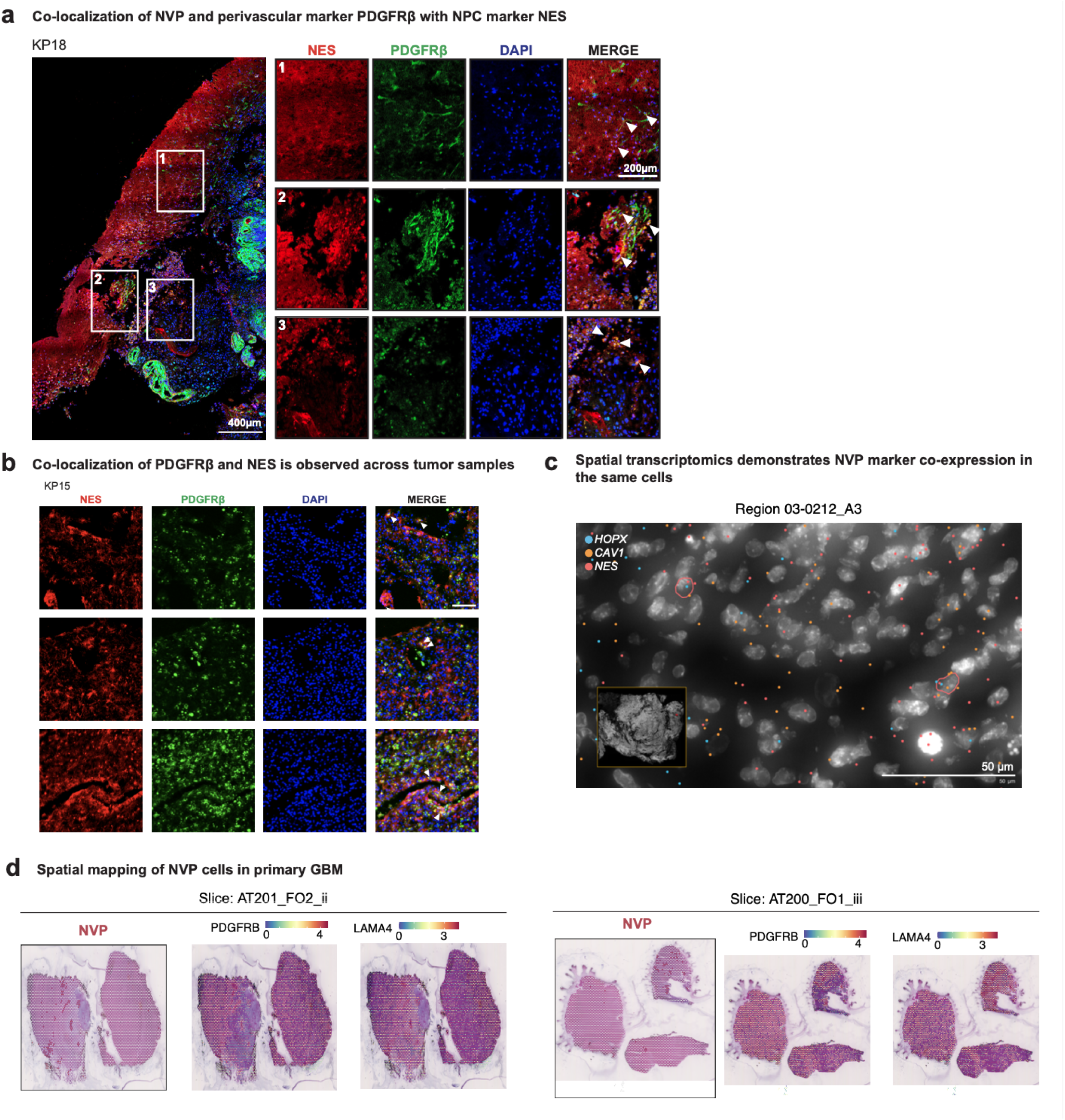
Co-localization of PDGFRβ and NES are observed surrounding vascular structures and in tumor parenchyma. a) In situ immunofluorescence was performed to assess co-localization of the NVP and perivascular marker PDGFRβ with the neural progenitor cell (NPC) marker NES in primary GBM tissue. Left: 20× tile scan from tumor KP18 showing co-localization of NES (red) and PDGFRβ (green); nuclei are counterstained with DAPI (blue). Insets 1-3 correspond to boxed regions in the overview image and display regions of marker co-expression, indicated by white arrows. Co-localization is observed both surrounding vascular structures and in tumor parenchyma. Scale bars: tile scans = 400 μm; inset panels = 200 μm (1.7X zoom). Representative images are shown. b) 20× imaging in additional tumor KP15 confirms the co-localization of PDGFRβ (green) and NES (red) with DAPI nuclear counterstain (blue). Representative fields demonstrate co-expression across tumor regions, with co-localization denoted by white arrows in merged images. Scale bar = 40 μm. Representative images are shown. c) Representative Xenium field demonstrating co-expression of NVP marker *CAV1* and neural progenitor markers *HOPX* and *NES* in primary GBM tissue. Transcript-level signals are overlaid on nuclear segmentation. Circled cells indicate triple-positive cells co-expressing NVP and neural progenitor markers. Scale bar, 50 µm. d) Representative Visium sections (AT201_FO2_ii and AT200_FO1_iii) annotated using RCTD-based label transfer from the GBM meta-atlas. Spots annotated as NVP are highlighted. Spatial feature plots for NVP markers *PDGFRB* and *LAMA4* are shown for each section (Methods).

**Supplementary Figure 4:**
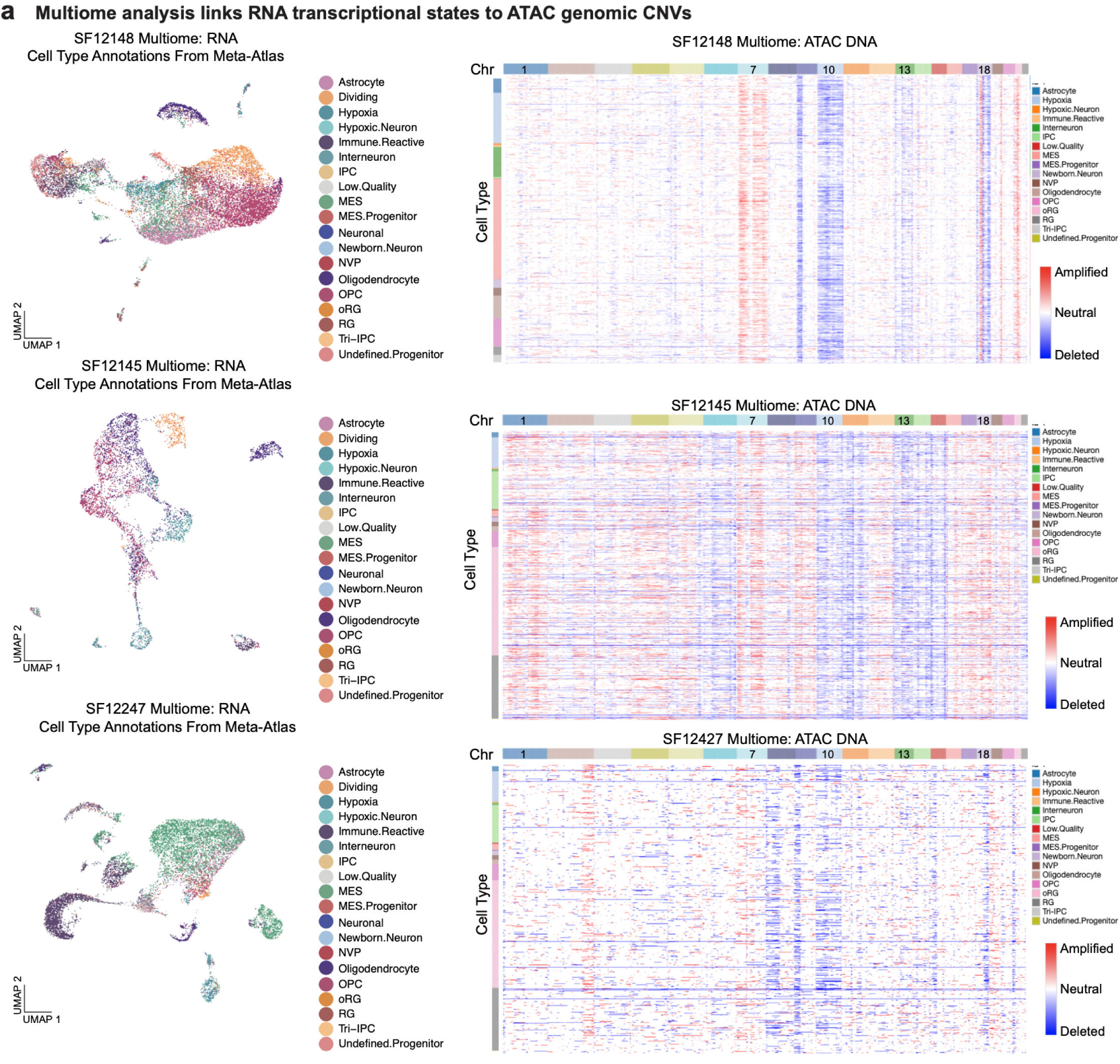
Multiome sequencing of GBM Identifies characteristic CNVs from ATAC-seq genomic regions across tumor cell types. a) Copy number variation (CNV) analysis was performed in joint scATAC-seq and scRNA-seq multiome profiling of 3 primary tumors. The scRNA data was used to identify cell types by using MapQuery to cross apply the cell types from our meta-atlas to the multiome. The UMAP of these cell types are shown for each tumor on the left, with each cell colored by cell type. The scATAC data was used to identify CNVs at the single-cell level with epiAneufinder^47^. Heatmaps^82^ of the results with the meta-data of the linked cell type identified from the RNA sequencing is shown on the right, with red indicating chromosomal amplification and blue indicating chromosomal deletion. Across all 3 tumors, CNVs were observed including the characteristic chr 7 amplification and chr 10 deletion, as well as other copy number variations. The heatmap is clustered by rows and shows the copy number variations are preserved across all cell types.

**Supplementary Figure 5.**
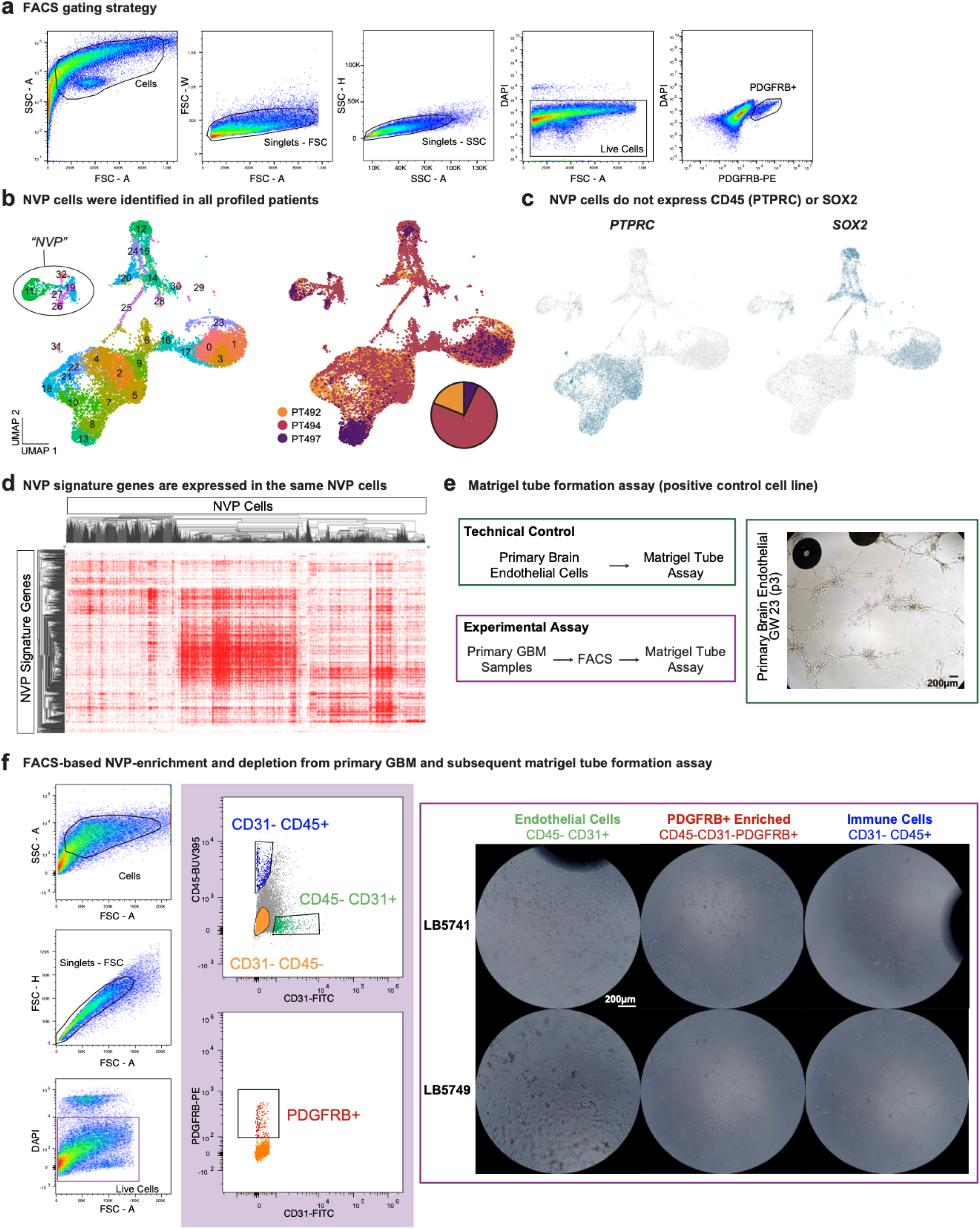
FACS enrichment allow for characterization of NVP. a) Fluorescence-activated cell sorting (FACS) gating strategy used to enrich for PDGFRβ+ cells. Cells were first gated by size (FSC-A vs. SSC-A), followed by exclusion of doublets using forward scatter (FSC-W vs. FSC-A) and side scatter (SSC-H vs. SSC-A). Live cells were selected via a DAPI-gate, and PDGFRβ+ cells were identified based on PE-conjugated PDGFRβ signal. b) UMAP visualization of sorted and whole tumor single-cell RNA-seq data. Left: Cells colored by cluster membership, revealing that NVP cells occupy a distinct region within the integrated dataset. Right: Cells colored by patient of origin (PT492, PT494, PT497), demonstrating that NVP cells are present across all profiled individuals. c) Feature plots showing minimal expression of canonical immune (CD45, encoded by PTPRC) and neural stem cell (SOX2) markers in the NVP population. d) Heatmap showing expression of NVP signature genes across NVP cells. The counts matrix includes only cells annotated as NVP in Figure 3b and genes included in the NVP signature. Genes and cells are hierarchically clustered, demonstrating coordinated expression of NVP signature genes within this defined population.

**Supplemental Figure 6.**
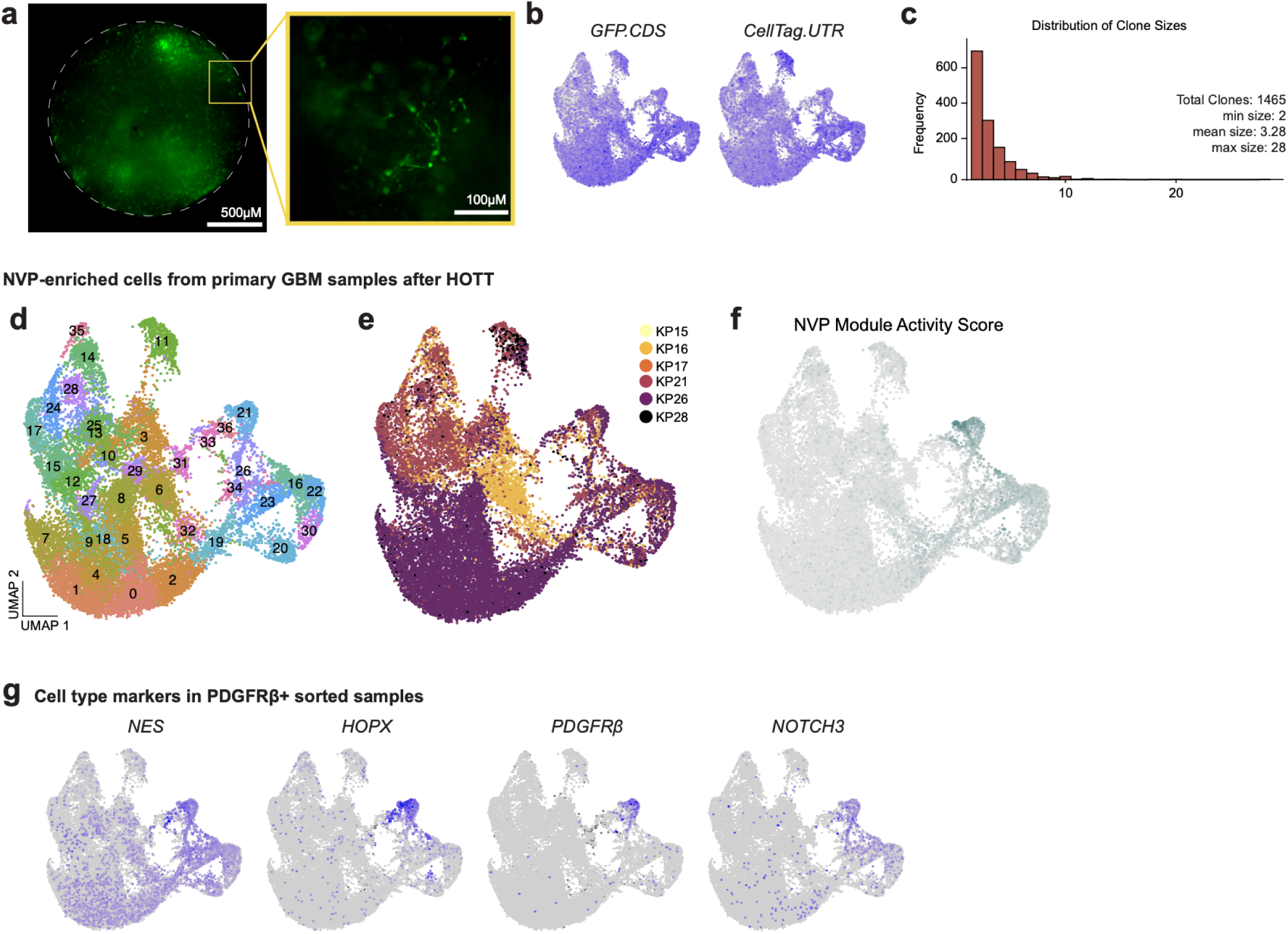
Clonal analysis of NVP-enriched human primary GBM cells following HOTT. a) Live-image of 3D cortical organoid with transplanted PDGFRβ+ CellTagged tumor cells in culture (4X magnification, scale bar = 500 μm). The inset depicts the cellular morphology of transplanted cells (20X magnification, scale bar = 100 μm). b) FeaturePlots depict the detection of CellTag associated genes GFP.CDS and CellTag.UTR, indicating that tumor cells were uniformly labeled by lentiviral transduction. c) Distribution of clone sizes. Total clones: 1465, minimum size: 2, mean size: 3.28, maximum size: 28. d) CellTagged PDGFRβ+ cells after dissociation from cortical organoid labeled by cluster e) Cells labeled by patient (n = 6) f) Cells were scored for the NVP signature derived from enriched NVP cells (Fig. 3, Methods), depicting high signature expression in the cluster annotated as “NVP”. g) FeaturePlots displaying the expression of cell type markers (NES, HOPX, PDGFRB, and NOTCH3) in PDGFRβ+ sorted, organoid-transplanted samples.

**Supplementary Figure 7.**
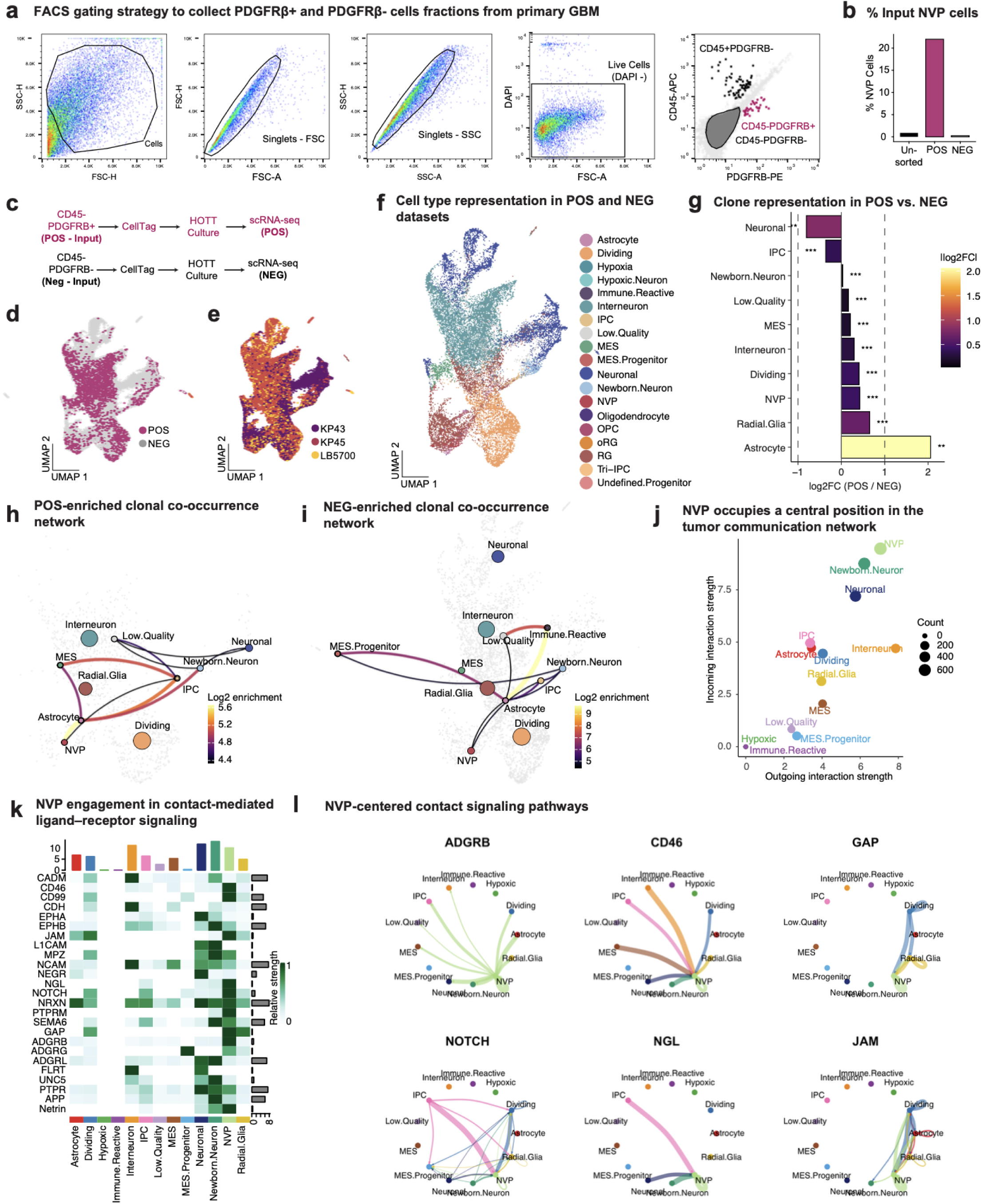
Depletion of PDGFRβ+ cells demonstrates signaling functions of NVP cells on other tumor cell populations. a) FACS gating strategy used to isolate PDGFRβ+ (POS) and PDGFRβ-(NEG) tumor cell populations. Cells were first gated by size (FSC-H vs. SSC-H), followed by sequential singlet selection based on forward scatter (FSC-A vs. FSC-H) and side scatter (SSC-A vs. SSC-H). Live cells were identified using a DAPI-gate. To eliminate immune cell contamination, cells were stained with CD45 and PDGFRβ antibodies. The CD45-PDGFRβ+ population was isolated as the enriched POS (magenta) fraction, while the CD45-PDGFRβ-population was collected as the depleted NEG (grey) fraction. b) After sorting, KP17 cells from the unsorted, POS (CD45-PDGFRβ+), and NEG (CD45-PDGFRβ-) fractions were directly captured for scRNA-seq prior to CellTag barcoding and HOTT culture. This allowed assessment of tumor composition prior to lineage tracing. Quantification of NVP cells in each fraction revealed strong enrichment in the POS condition, where NVP cells accounted for approximately 20% of the total population, compared to ∼1% in the unsorted tumor and <1% in the NEG fraction. c) Lineage tracing of PDGFRβ+ (POS) and PDGFRβ-(NEG) tumor cell populations (n = 3 tumors) using CellTag barcoding and orthotopic human organoid tumor transplantation (HOTT). Following FACS isolation, cells from each fraction were infected with a CellTag lentiviral barcode library (Addgene), cultured in the HOTT system for two weeks, and subsequently harvested for scRNA-seq (n = 22,321 cells). d) Integrated UMAP showing POS and NEG lineage-traced cells, colored by input condition. e) UMAP showing the same cells colored by patient of origin. f) Integrated UMAP annotated by cell type identity, derived via mapping to the CellTagging^19^ dataset. g) Weighted log₂ fold change in relative clone member cell-type abundance (n = 1592 clones), computed as log₂(POS / NEG) at the cell level (weights = 1/p(CellType|Condition)). Values are capped at ±3 for visualization. Dashed lines indicate ±1 (two-fold change). Asterisks denote Benjamini–Hochberg FDR-adjusted significance (* <0.05; ** <0.01; *** <0.001). Cell types with <10 cells were excluded. h) Clone co-occurrence network for clones enriched in the POS condition. Edges connect cell-type pairs that co-occur within inferred clones more frequently than expected by chance (log₂ observed/expected). Edge thickness and color indicate enrichment magnitude. Node size reflects overall cell-type abundance. i) Clone co-occurrence network for clones enriched in the NEG condition, shown as in (h). NVP-depleted clone networks reflect increased reliance on communications centered around other progenitors, namely MES.Progenitors, Radial Glia, and IPCs. j) NVP cells function as signaling hubs with context-dependent interaction profiles. Cell-cell communication analysis was performed using ligand-receptor interaction modeling across all lineage-traced cells. Left: Scatter plot summarizing global signaling activity for each cell type, aggregated across both PDGFRβ+ (POS) and PDGFRβ-(NEG) conditions. The x-axis represents outgoing interaction strength (signaling sent), and the y-axis represents incoming interaction strength (signaling received). k) Heatmap depicting NVP participation in receptor-ligand interactions involved in cell-cell contact. Rows represent contact-associated ligand-receptor pairs, and columns represent annotated tumor cell types. Signal intensity reflects the relative strength of predicted interactions based on transcriptomic expression. NVP cells display broad engagement across multiple receptor-ligand families, highlighting their role as a key mediator of contact-dependent intercellular communication within the tumor. l) Chord diagrams displaying cell-cell contact signaling interactions. Druggable, contact-mediated pathways shown include ADGRB, CD46, GAP, NOTCH, NGL, and JAM with NVP cells serving as signaling sources to multiple recipient states.

**Supplementary Figure 8.**
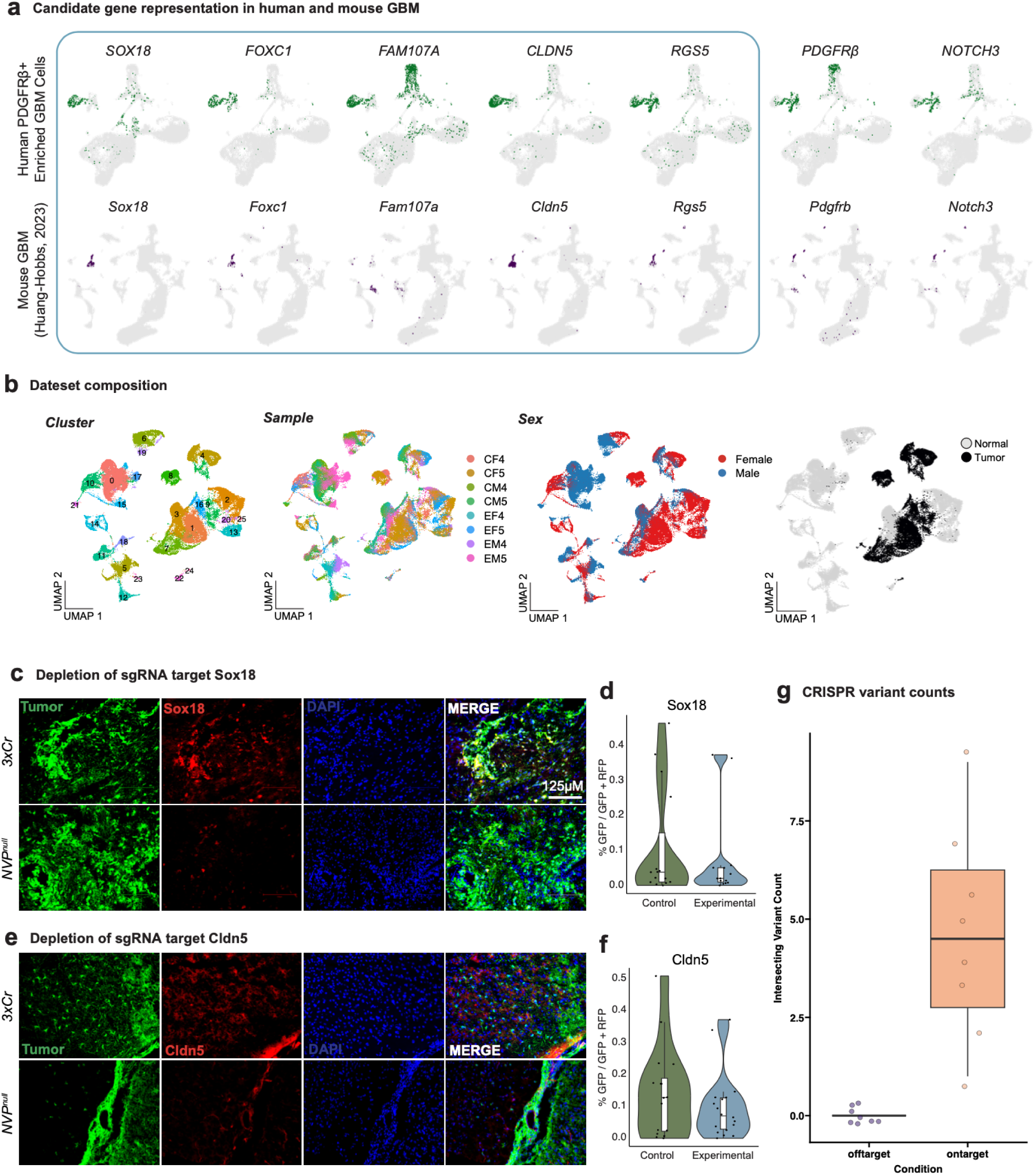
Candidate selection for *in vivo* ablation was based on specificity to human and mouse NVP. a) Expression of candidate genes in human and mouse GBM. UMAP plots show gene expression in PDGFRβ+ enriched human GBM cells (top row) and in a mouse GBM model (bottom row, data from mouse GBM cells36 (bottom row). Candidate genes were selected based on specificity to the annotated “NVP” cluster in the human and mouse datasets. Genes targeted in CRISPR-Cas9 ablation experiments are outlined in blue. PDGFRβ and NOTCH3 are included as reference perivascular markers. b) Dataset composition for snRNA-seq analysis of CRISPR-targeted tumors. UMAP plots depict clustering by transcriptional identity, experimental sample, biological sex, and the tumor vs. nomral annotation. Data are derived from *3×Cr* control and *NVP^null^* tumors. c) Validation of Sox18 depletion following *in vivo* CRISPR targeting. Immunofluorescence was performed on tumors collected at postnatal day 70 (P70), the same time point as transcriptomic profiling, to assess reduction of Sox18 protein in response to sgRNA targeting. Representative 20X images are shown for control (*3xCr*) and experimental (*NVP^null^*) tumors. Tumors from n = 3 mice per condition were analyzed. Sections were stained for GFP+ tumor cells (green), Sox18 (red), and DAPI (blue); merged panels display co-localization. d) Quantification of Sox18 protein expression using Synbot, shown as the fraction of GFP+ cells co-expressing Sox18, demonstrates reduced Sox18 in experimental tumors compared to controls (n=15 images per condition). e) Validation of Cldn5 depletion following *in vivo* CRISPR targeting. Immunofluorescence was performed on tumors collected at postnatal day 70 (P70), concurrent with transcriptomic profiling, to assess reduction of Cldn5 protein following sgRNA targeting. Representative 20X images are shown for control (3xCr) and experimental (NVP^null^) tumors. Tumors from n = 3 mice per condition were analyzed. Sections were stained for GFP+ tumor cells (green), Cldn5 (red), and DAPI (blue); merged panels display co-localization. f) Quantification of Cldn5 protein expression using Synbot, shown as the fraction of GFP+ cells co-expressing Cldn5, demonstrates reduced Cldn5 in experimental tumors compared to controls (n=15 images per condition). g) Exome sequencing reveals on-target but not off-target CRISPR-Cas9 activity in NVP^null^ tumors. Whole-exome sequencing was performed on DNA isolated from control and experimental tumors collected at p70 and at terminal disease. Variant analysis was conducted using Mutect2, with control tumors used as reference samples. On-target sites were defined as predicted cut sites identified by CRISPOR, and the top five highest-scoring off-target loci per sgRNA were included for evaluation. The intersecting variant count for each locus is shown for on-target versus off-target sites across n=4 experimental tumors from each timepoint (total n=8). On-target mutations were observed, whereas off-target sites showed no detectable editing activity.

